# Faroese Whole Genomes Provide Insight into Ancestry and Recent Selection

**DOI:** 10.1101/2025.05.20.655212

**Authors:** Iman Hamid, Ólavur Mortensen, Alba Refoyo-Martínez, Leivur N. Lydersen, Anne-Katrin Emde, Melissa Hendershott, Katrin D. Apol, Guðrið Andorsdóttir, Jonas Meisner, Kaja A. Wasik, Fernando Racimo, Stephane E. Castel, Noomi O. Gregersen

## Abstract

The Faroe Islands are home to descendants of a North Atlantic founder population with a unique history shaped by both migration and periods of relative isolation. Here, we investigate the genetic diversity, population structure, and demographic history of the islands by analyzing whole genome sequencing data from 40 participants in the Faroe Genome Project. This represents the first whole genome sequencing panel of this size from the Faroe Islands. We observed numerous putatively functional private alleles, including stop gain variants and high impact missense variants in the cohort. Faroese individuals had a higher proportion of their genomes contained in long runs of homozygosity than other European groups, including Finnish, suggesting a more recent or stronger bottleneck in the Faroese population. Signals of positive selection were identified at loci containing genes that play roles in vitamin D and dietary fat absorption and DNA repair, while increased diversity on lactase persistence haplotypes was observed. Fine-scale analysis of haplotype structure in present-day and ancient European genomes revealed genetic affinities with ancient Iron Age individuals from the North and West of Europe, providing evidence for potential contributions to the Faroese gene pool from Celtic and Viking populations as well as information about the temporal order in which these events happened. This study highlights the impact of evolutionary processes, such as ancient admixture, founder events, and positive selection, on the present-day genetic architecture of North Atlantic founder populations like the Faroe Islands.

## Introduction

The Faroe Islands, nestled in the North Atlantic Ocean between Iceland and Norway, are home to the descendants of a North Atlantic founder population with a rich cultural heritage and a unique history shaped by both migration and periods of relative isolation. The exact settlement history of the islands is unclear, though historical records and analysis of Y-chromosome microsatellite markers point to a few founders most likely having arrived primarily from Scandinavia and the British Isles starting around the 9th century C.E.^1–3^ However, archeological evidence supports a possible earlier settlement of the island around the 4th-6th centuries C.E.^4^ Studies of mtDNA reveal an excess of maternal ancestry from the British Isles, while Y chromosome studies reveal an excess of paternal ancestry from Scandinavia, suggesting sex-biased admixture between these ancestral groups during the founding of the population.^2,5^ Since settlement and early waves of migration, the Faroe Islands have been mostly isolated and have experienced minimal immigration and population growth until recent years, with a census size of about 4,000 people in the 1700s^6^ increasing to over 54,000 as of August 2023 (https://hagstova.fo/en/population/population/population). Patterns of genetic diversity and linkage disequilibrium from a few genetic markers suggest a founder event, followed by a historically small population size and subsequent rapid expansion.^7^

Overall, despite the small size and remote location of the Faroe Islands, the above evidence suggests that the genetic makeup of Faroese people may have been influenced by waves of early migration and admixture from various northwest European and Scandinavian populations. However, no population genomic studies have yet been carried out on whole–genome sequencing data to date, with the exception of one study investigating relatedness and autozygosity for a limited sample size of eight individuals.^8^ An in-depth analysis of the genomic architecture of the Faroese may reveal how the islands’ demographic history has contributed to present-day health and disease in this population. This is of particular interest as the Faroese have a high burden of certain diseases relative to global and other European populations, such as inflammatory bowel disease and type 2 diabetes, among others.^9–18^ The Faroe Genome (FarGen) Project set out to understand how the genetic diversity of the Faroese contributes to health.^19^

Moreover, studying the genetic diversity of the Faroese provides potential insight into human migration, adaptation, and population structure in the North Atlantic region. Recently developed haplotype-based methods can provide fine resolution for inferring shared ancestry among individuals and the detection of population-specific haplotypes, but they require panels of whole-genome sequencing data.^20^ These methods serve to better account for recent patterns of population structure and cryptic relatedness in population-based genetic studies, particularly in populations like the Faroe Islands with strong founder effects.

In this study, we present the first whole genomes sequenced as part of the FarGen Project. A set of 40 individuals were selected to optimally represent the genetic diversity of the broader Faroese population, with the aid of genealogical data reaching back to approximately 1650 CE. Whole genome sequencing (WGS) data was generated and used to identify putatively functional alleles enriched in the Faroese population, assess ancestry patterns within contemporary genomes, map signals of recent positive selection, and analyze local ancestry in a combined dataset of ancient and contemporary genomes of European descent spanning a period of 3,000 years.

This study provides insight into the genetic variation, demographic history, and selection landscape of the Faroese population. These first whole genomes from FarGen may serve as a useful reference for studies on the broad implications of various evolutionary genetic processes, including bottlenecks, ancient admixture, and positive selection. More in-depth studies may expand this current study by further investigating the genetic architecture of the Faroese to unravel the demographic and evolutionary history of the population and its impact on complex traits and diseases in the islands.

## Results

### Whole Genome Sequencing of Faroese Individuals

The Faroese Multi Generation Register (https://fargen.fo/research/multi-generation-registry) was used to reconstruct a single connected genealogical tree for the 1,541 participants in the first phase of the FarGen cohort.^19,21^ Individuals with fewer than six direct ancestors (two parents and four grandparents) recorded in the registry were excluded. Pairwise kinship coefficients were estimated from the genealogical tree and used to perform relatedness pruning with a threshold of 2^−6^ resulting in 332 minimally related individuals. We defined six geographical regions of the Faroe Islands using language dialects, and assigned individuals to these regions based on their place of birth (Fig. 1A). A total of 40 minimally related individuals were selected for inclusion in the study, with five to eight individuals sampled from each region.

**Figure 1.**
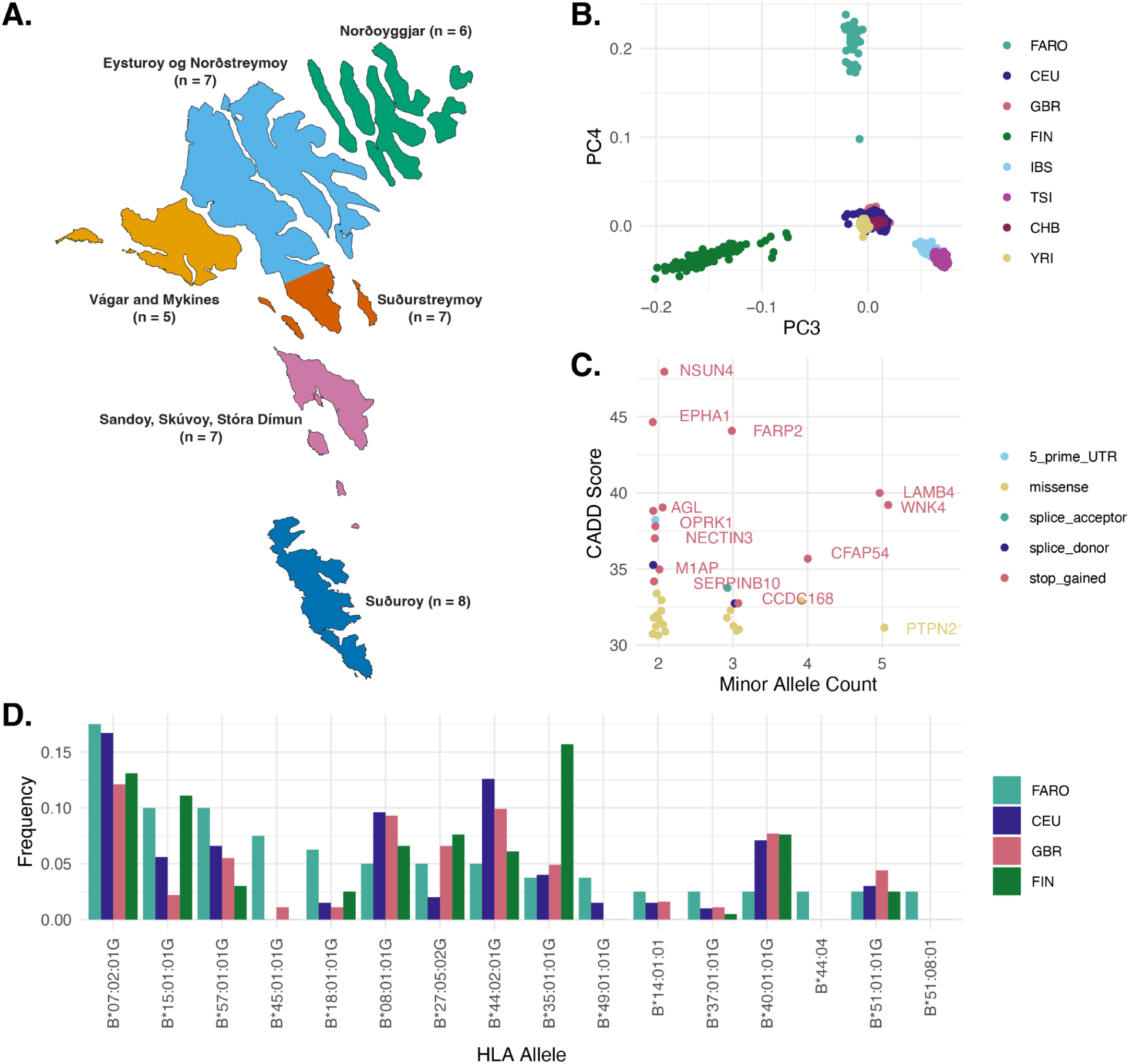
A Faroese whole genome reference. **A)** Map of the Faroe Islands, colored by the six sampling regions. The number of minimally related FarGen participants from each region selected for whole genome sequencing is indicated. **B)** Principal component analysis (PCA) of Faroese genomes jointly called with relevant 10000 Genomes reference data shows separation of European groups by PCs 3 and 4 (FARO, Faroese, CEU, Central Europeans, GBR, British, FIN, Finnish IBS, Iberian, TSI, Tuscan, CHB, Han Chinese, YRI, Yoruban). **C)** Faroese enriched putatively functional alleles visualized by minor allele count, CADD score, and Variant Effect Predictor (VEP) consequence. Variants shown are those with CADD > 30 and at least two minor alleles observed in Faroese individuals, and no minor alleles observed in Finnish or Northern European reference individuals. **D)** HLA-B allele frequencies for alleles detected at least twice in Faroese individuals. In this cohort, 1 minor allele corresponds to an allele frequency of 1.25%.

The genomes of the 40 individuals were sequenced to a median depth of 20x in the Faroe Islands at the FarGen laboratory, and they all passed quality control metrics (Fig. S1, Table S1). Variant calling was performed jointly with relevant reference genome panels, including 1000 Genomes high-coverage data from Northern European (CEU & GBR, N = 190), Southern European (TSI & IBS, N = 214), Finnish (FIN, N = 99), East Asian (CHB, N = 103), and West African (YRI, N = 108) individuals, and imputation was performed within the cohort using an approach we have previously described.^22,23^ The first component of principal component analysis (PCA) on the jointly called genotype data captured the cline between West African (YOR) and all other individuals, and the second component captured the cline between East Asian (CHB) and all other individuals (Fig. S1). Principal components three and four separated European individuals, with Faroese individuals forming a distinct cluster from Finnish, Northern European, and Southern European reference groups (Fig. 1B).

Variant calls were annotated with predicted functional impact and allele frequency across the reference groups. Using these annotations, we identified 35 putatively functional alleles present in our Faroese panel, that are unobserved in the European mainland reference panels included in this study (CADD > 30 and at least two minor alleles observed in Faroese individuals, and no minor alleles observed in Finnish or Northern European reference individuals, Table S2). These included 13 stop gain variants and 18 missense variants, and a maximum minor allele count of 5, corresponding to a frequency of 6.25% in the cohort (Fig. 1C).

### HLA-B Allele Frequencies

Observational evidence from the FarGen project recruitment data suggest that ankylosing spondylitis (AS) may be at a higher prevalence in the Faroe Islands ; however, more formal epidemiological studies are required to confirm this observation. The major histocompatibility complex (MHC) plays a role in various autoimmune diseases that may be at higher prevalence in the Faroes, including ankylosing spondylitis and other more common diseases like inflammatory bowel disease.^24,25^ In particular, HLA-B*27 is associated with ankylosing spondylitis (AS), with approximately 80-90% of AS patients carrying the HLA-B*27 allele.^25^ It explains about 30% of the heritability, and ∼6-8% of European populations are carriers of HLA-B*27.^25–28^ Using the WGS data, we genotyped human leukocyte antigen (HLA) alleles with HLA*LA.^29^ We provide HLA-B allele counts and allele frequencies in the Faroese cohort as well as the allele frequencies in 1000 Genomes British (GBR), Central European (CEU), and Finnish (FIN) individuals (Table S3). To the best of our knowledge, there have not been any larger studies of HLA-B allele frequencies in Faroese individuals, and none are currently recorded in the Allele Frequency Net Database.^30^

The most frequent HLA-B allele in the Faroese cohort is HLA-B*07:02 (17.5% of haplotypes), which is a common haplotype in European populations (Fig. 1D).^31,32^ We did not observe a substantial difference in HLA-B*27 allele frequency in Faroese individuals (6.25% with four B*27:05 and one B*27:01 calls) as compared to other European reference groups (2-7.6%). While 80-90% of people of European ancestry with AS carry the HLA-B*27 allele, only ∼6% of HLA-B*27 carriers in the US and Europe have AS.^33–35^ The low frequency of the HLA-B*27 allele in this Faroese cohort in the population broadly suggests that if AS is at a higher prevalence, there may be other underlying genetic or environmental factors that explain some of the increased risk.

### Population Structure and Relatedness

Pairwise kinship was calculated in the cohort using *popkin*.^36–38^ Clustering by geography was observed when including global reference populations in the kinship calculation. The Faroe Islands have high pairwise kinship within the cohort when compared to other global populations, which may be indicative of recent bottlenecks (Fig. S2A). The Faroese individuals do not show obvious clustering by region, though this is expected given the relatedness pruning during sample selection (Fig. S2B).

We also looked at runs of homozygosity (ROHs) in the Faroese and reference cohorts (Fig. 2), which can provide further insights into the demographic history of the population. As the Faroese population likely experienced a founder event during the settlement of the islands followed by rapid population size expansion in recent generations, we would expect to see more of the genome contained in ROHs compared to other global populations that have not experienced as strong a bottleneck.^39^ When looking at the sum total amount of the genome in ROHs, we found overall elevated levels of ROH in the European and Asian groups included in this analysis, most likely reflecting ancestral out-of-Africa bottlenecks for Eurasian populations (Fig. 2, top panel). However, the Faroese population did not have an elevated total amount of the genome contained in ROHs compared to other European groups.

**Figure 2.**
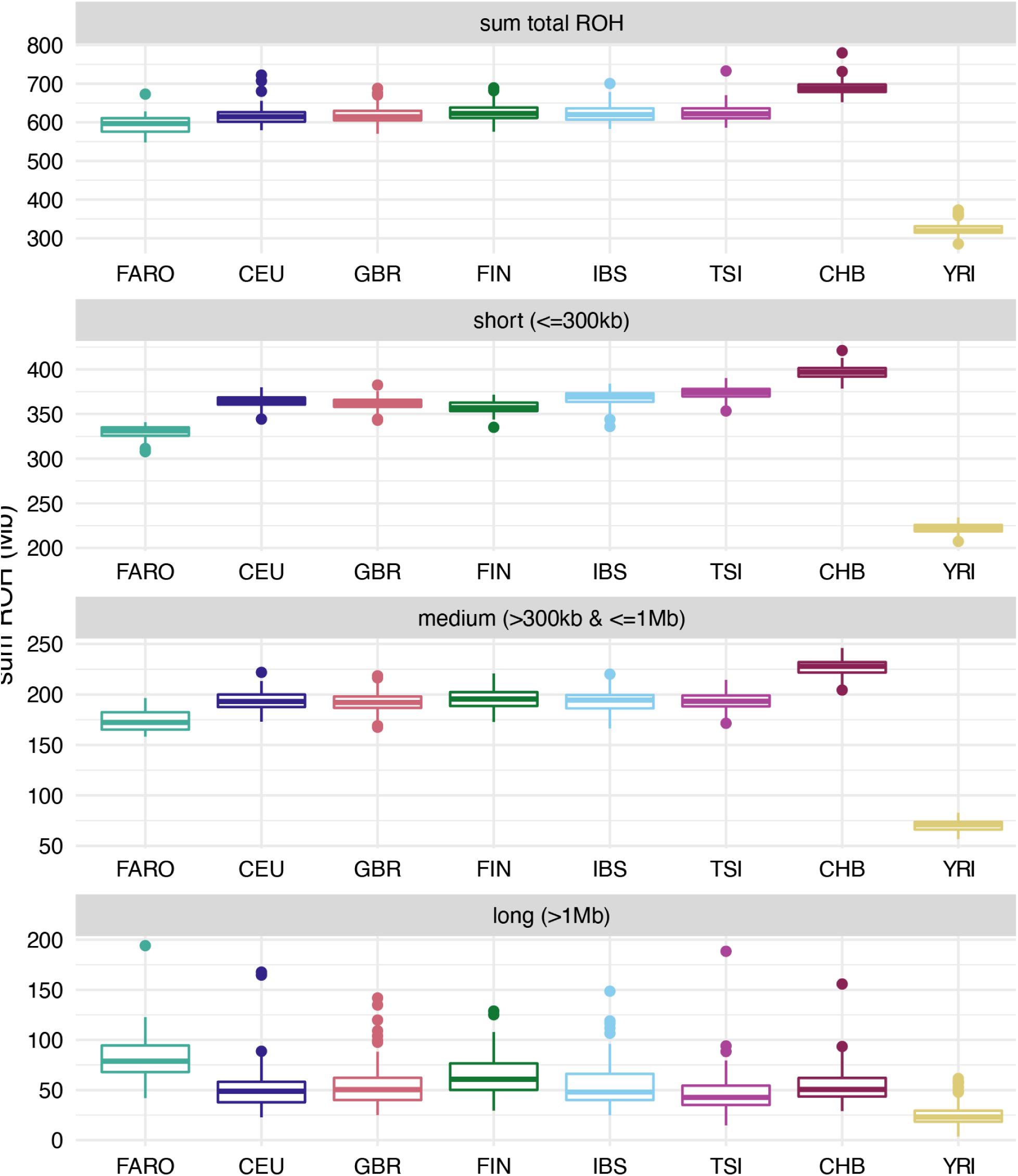
Runs of homozygosity by group. Amount of the genome (Mb) contained in runs of homozygosity (ROH) stratified by group. Top panel is the sum total of the genome contained within ROH, with the other panels showing this split by length (short, medium, and long).

To explore this further, we calculated the sum total amounts of ROH at different size categories (small, medium, large) (Fig. 2, bottom 3 panels). We see that, on average, the Faroese individuals have less of the genome contained in short (<300 Mb) and medium ( > 300 Kb and <= 1 Mb) ROHs compared to other European groups, but more of the genome contained in long ( > 1Mb) ROHs. Short and medium ROH are chunks inherited from older ancestors and reflect older events, for example, an ancient population bottleneck or founder event that has resulted in lower overall haplotype diversity, yet with enough time for recombination to break up haplotypes, while long ROH reflects chunks inherited from recent ancestors and can reflect more recent bottleneck events.^39^ Interestingly, the average amount of an individual’s genome that is contained in ROHs extending over 1Mb in length is higher in the Faroese population (∼82.5 Mb) than the Finnish reference individuals (∼63.9 Mb) and any of the other groups analyzed. Additionally, distribution in ROH lengths across all individuals stratified by group shows that, on average, there is a higher proportion of long ROH particularly in the 5-15 Mb range in the Faroese cohort relative to the other cohorts (Fig. S3). This is consistent with a more recent or stronger bottleneck or founder event.

### Signals of Positive Selection

We investigated signals of recent or ongoing positive selection in this Faroese cohort by calculating both the integrated haplotype score (iHS)^40^ and cross-population expected haplotype homozygosity (XPEHH) using *hapbin*.^41^ The sample size of the WGS cohort is relatively small (n=40), so our ability to detect signals of selection is limited. For comparison, we also calculated iHS in British individuals from the 1000 Genomes (GBR) that were included in joint calling and passed subject-level filters (n=90). The iHS values were standardized genome-wide and two-tailed p-values were computed according to the standard normal distribution. We additionally calculated q-values (the minimum False Discovery Rate (FDR) should a test be considered significant) for each test,^42^ and determined p-value significance thresholds at which FDR < 0.01 and FDR < 0.001 for each population (see Methods). We observed that a number of significant selection signals were shared between the Faroese and British cohorts, which is not unexpected given the relationship between these two populations (Fig. 3A-B). The strength of these signals did differ from one population to the other, though this may be due to differences in sample size or changes in selection pressure after the populations diverged. To better identify population-specific signals, we also calculated XPEHH comparing Faroese and British haplotypes, and identified significance cutoffs following the same approach described above (Fig. 3C).^43^ Any extreme positive values of this statistic indicate longer haplotypes at a focal marker in the Faroese cohort compared to the British cohort, while extreme negative values indicate the reverse. Therefore, positive values are indicative of selection signals in the Faroese cohort. Across both tests, we highlight 20 loci with the most extreme values for these statistics, serving as evidence of positive selection in the Faroese genomes at those loci (Table S4).

**Figure 3.**
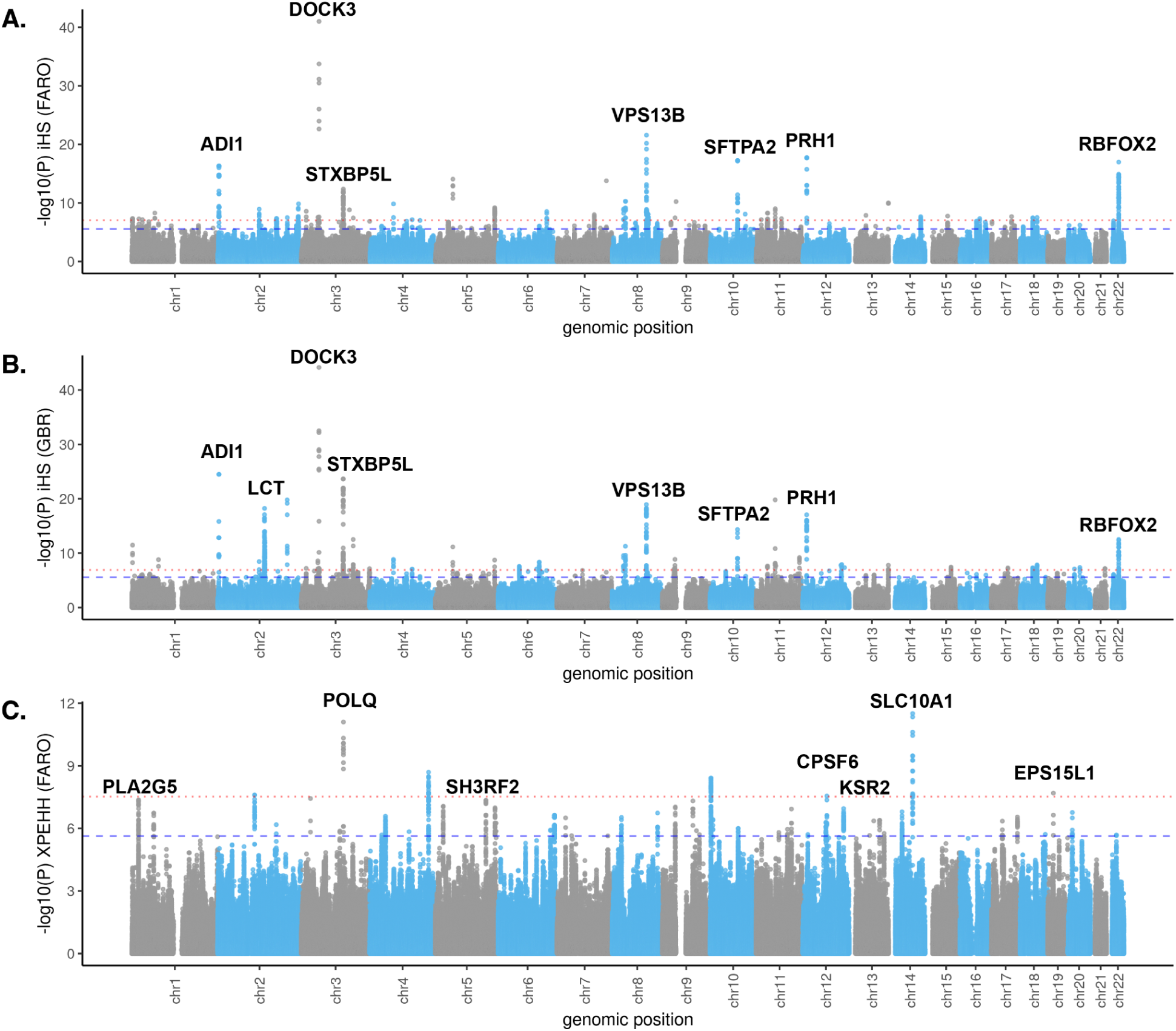
Selection scan results for Faroese and British cohorts. **A)** Log transformed two-tailed p-value of the standardized integrated haplotype score (iHS) in the 40 Faroese genomes (FARO). **B)** Log transformed two-tailed p-value of the standardized iHS for 90 British WGS samples from 1000 Genomes (GBR). **C)** log transformed two-tailed p-value for the standardized cross-population expected haplotype homozygosity (XPEHH) for FARO vs GBR (only positive values, which indicate selection in FARO, are plotted). Some genes in the top loci are indicated on each plot. The p-value cutoffs which correspond to a False Discovery Rate (FDR) at 0.01 and 0.001 are respectively indicated by the red dotted line and blue dashed line in each plot. **A)** For iHS in FARO, these cutoffs are p = 2.72 x 10^-6^ (FDR = 0.01) and p = 9.20 x 10^-8^ (FDR = 0.001). **B)** For iHS in GBR, the cutoffs are p = 2.78 x 10^-6^ (FDR = 0.01) and p = 1.75 x 10^-7^ (FDR = 0.001). **C)** For XPEHH in FARO vs GBR, the cutoffs are p = 2.35 x 10^-6^ (FDR = 0.01) and p = 3.01 x 10^-8^ (FDR = 0.001). See Methods for details on p-value and FDR estimation.

One signal that has been consistently observed across northern European populations is in the *LCT*/*MCM6* region, corresponding to positive selection for lactase persistence alleles.^40,43–45^ Interestingly, this region showed strong iHS signals (|standardized iHS| > 8) in GBR and was considered genome-wide significant in our analysis (minimum p = 5.95 x 10^-19^, q = 7.82 x 10^-14^) (Fig. 3B), but the signal is weaker in the Faroese (|standardized iHS| > 4) and was not considered significant in our analysis (minimum p = 3.57 x 10^-6^, q = 0.0118) (Fig. 3A). To investigate the haplotype structure further, we plotted the decay in expected haplotype homozygosity (EHH)^46^ and haplotype furcation around one of the lactase persistence alleles (rs4988235; chr2_135851076_G_A) for the Faroese and British cohorts using the *rehh* package (Fig. 4A-D).^47^ The decay and furcation plots are centered around the focal marker, and a furcation occurs when unique haplotypes arise at an allele, similar to a tree splitting into branches. Thicker branches in the furcation plot indicate higher frequency of that haplotype in the population. Significant differences between the furcation patterns for an “ancestral” (reference) and “derived” (alternate) allele correspond to extreme iHS values and therefore are indicative of strong positive selection. For an alternative view of the region, we used *Haplostrips* to visualize the haplotype structure from chr2:135677850-135986443 (Fig. 4E).^48^ From these plots, we observed far less diversity on the lactase persistence haplotype in GBR, consistent with a stronger selection signal. This may be explained by shared selection on the ancestral northern European branch followed by either relaxed selection for lactase persistence or population-specific drift in the Faroes after the population split from other northern European groups and settled the archipelago.

**Figure 4.**
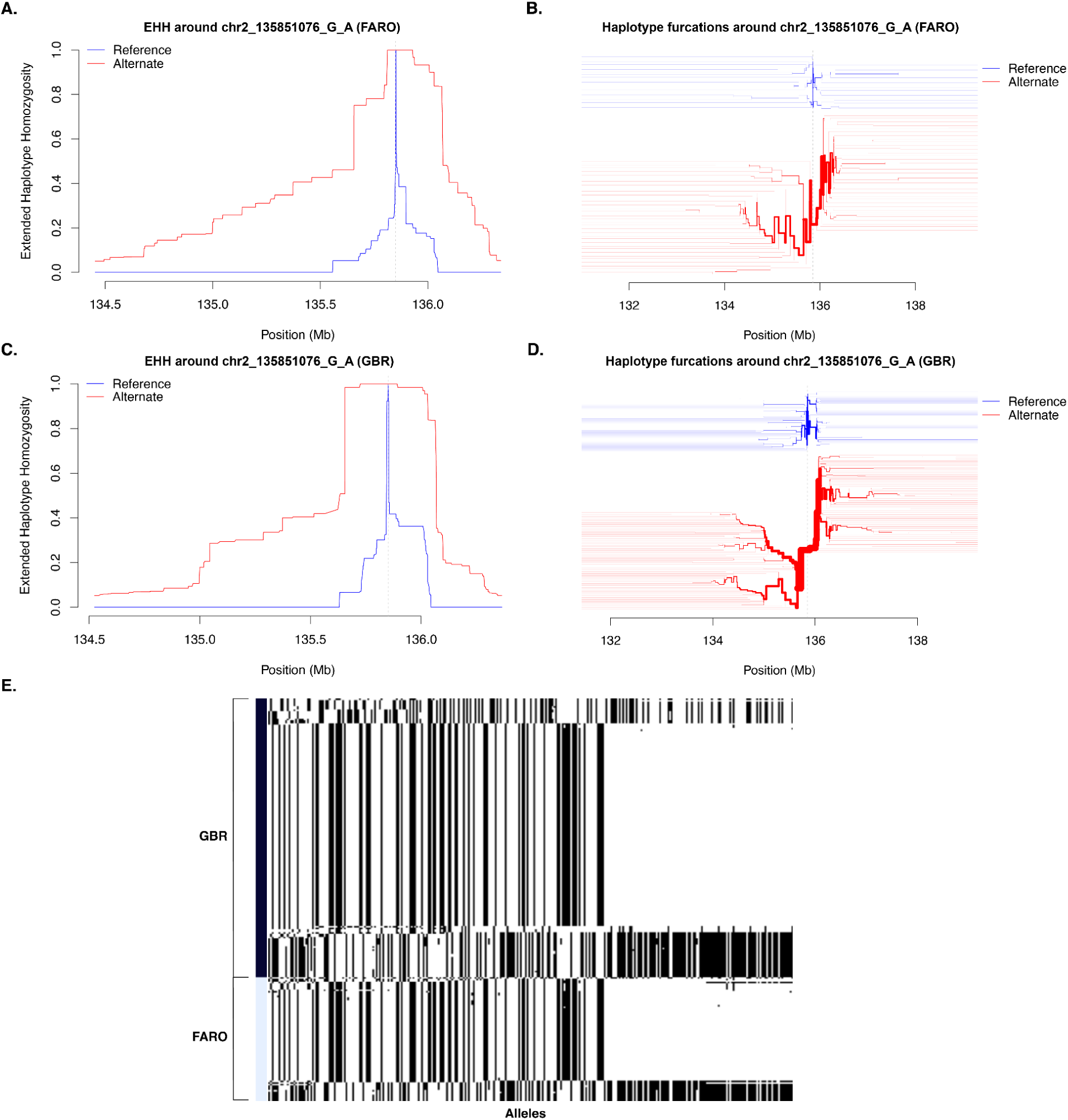
Haplotype visualizations for the LCT/MCM6 locus. **A)** Decay in Expected Haplotype Homozygosity (EHH) and **B)** haplotype furcation plot for FARO centered on lactase persistence allele rs4988235; chr2_135851076_G_A. **C)** Decay in EHH for GBR and **D)** haplotype furcation for GBR centered on the same allele. **E)** *Haplostrips* visualization of haplotype structure in the region chr2:135677850-135986443. In this panel, columns correspond to segregating alleles, and rows correspond to individuals. In the haplotype furcation plots (panels **B** & **D**), the haplotypes for the reference allele (G) are in blue, and those for the alternate allele (A) are in red.

One of the top XPEHH signals in the Faroese WGS cohort included variants in *SLC10A1* (Fig. S4), a sodium/bile acid transporter that plays a role in circulating bile salts to and from the liver and small intestine for the absorption of dietary fat and fat-soluble vitamins such as vitamin D.^49–52^ SLC10A1 deficiency has been associated with familial hypercholanemia, or elevated concentrations of bile acids, which can lead to fat malabsorption and vitamin D deficiencies among other secondary health conditions.^50,53,54^ Another top XPEHH signal included variants in *POLQ*, encoding for a DNA polymerase which plays a role in DNA repair (Fig. S5).^55–59^ *POLQ* has been shown to be involved in various cancers in mice and humans, in particular skin, stomach, lung, breast, and colon cancers.^55,59–62^

### Fine-Scale Structure and Connections to Ancient Genomes

Given that early Faroese settlers have documented historical relations to both Northern European Vikings and Northwestern European Celtic communities, we sought to study fine-scale genome-wide ancestry relationships between the sequenced Faroese genomes and publicly available ancient genomes from Iron Age and Viking Age Europe. We downloaded 616 ancient imputed genomes from Allentoft *et al.* 2024, spanning from the present-day to the late Bronze Age from Europe and focusing specifically on West- and North-Europe, including ancient Faroese genomes.^63^ We incorporated these genomes into a combined panel including our present-day Faroese dataset and used the software HaploNet to infer fine-scale population structure based on patterns of haplotype similarity across the genome.^20^

HaploNet identified five ancient sources, through unsupervised ancestry estimation. We then used HaploNet to model both ancient and present-day Faroese genomes as composites of any of the five ancestries through supervised ancestry estimation. We used the ancestral haplotype cluster frequency estimates from individuals not found in the Faroe Islands in order to estimate the admixture proportions of these sources in the European mainland. We used these admixture proportions to label the sources based on the locations in the map where these ancestries tend to be maximized in the Iron Age and Viking Age periods. The resulting labels were: “Steppe”, “East Europe”, “Levant and East Mediterranean”, “West Europe” and “North Europe”. For example, the ancestry labeled “West Europe” is maximized in individuals predominantly found in Celtic contexts (e.g. Roman and Iron Age Britain, Iron Age France) while the ancestry “North Europe” is maximized in individuals characteristic of historically Viking or pre-Viking contexts (e.g. Iron Age individuals from Denmark, as well as Viking Age individuals from Denmark, Norway, Sweden, and Estonia). However, we note there is no one-to-one correspondence between archaeological context and genetically inferred ancestry, and that many mainland individuals contain inferred ancestries from diverse sources. Indeed, Margaryan *et al*. 2020 showed that Viking-context individuals can derive ancestries from multiple Bronze and Iron Age sources across Europe.^64^

We then focused on the frequency of the ancestry sources in the Faroese individuals. We find that the present-day Faroese individuals are predominantly composed of roughly equal proportions of “West” and “North Europe” ancestry, while “East” and “South Europe” ancestries are detected at much lower frequencies (40% North Europe, 33.1% West Europe, 12.2% Levant, 8.5% East Europe, 6.2% Steppe). Present-day ancestry proportions are nearly identical to those found in the Faroese ancient samples from the Sandur church site in Sandoy and dated to the 17th or 18th centuries based on their archaeological contexts (37.3% North Europe, 35.6% West Europe, 12.0% Levant, 8.1% East Europe, 7.0% Steppe) (Fig. S6).^64^ Margaryan *et al.* also sequenced a Faroese sample that was excavated from the á Bønhúsfløtu site in the village of Hvalbøur on Suðuroy, and was contextually-dated to be approximately 800 years old. This individual is inferred to be almost entirely composed of “West Europe” ancestry (Fig. S7).

We utilized haplotype cluster likelihoods to explore population structure in the same set of samples. When plotting ancient European samples together with the Faroese samples (Fig. 5), we observe that present-day Faroese individuals (circled in black) separate from the ancient Europeans along the second principal component (PC2), as do the older Faroese samples from the 17th and 18th centuries (circled in red). This perhaps suggests a bottleneck process that differentiates the 17th/18th-century and present-day Faroese from the rest of the ancient European samples, in concordance with the above ROH results. Notably, the 800-year-old sample of a Faroese individual with predominantly “West Europe” ancestry does not fall along the Faroese PC2 cline, suggesting that this individual might predate the bottleneck.

**Figure 5.**
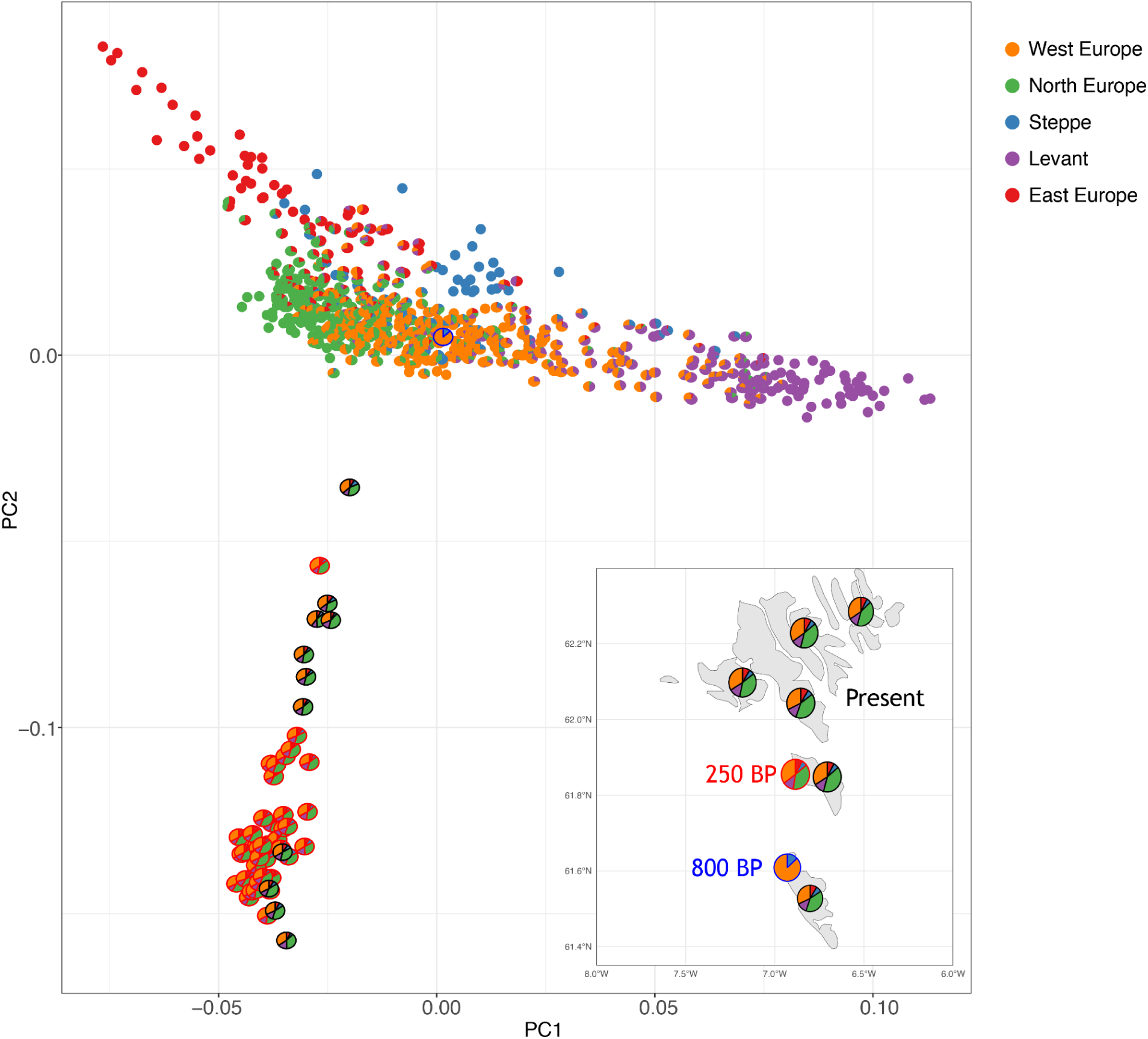
PCA analysis of 616 ancient imputed genomes from Europe and 40 present-day Faroese genomes. Each individual is depicted as a pie chart, showing ancestry proportions estimated using Haplonet. Ancestry proportions for ancient individuals were estimated unsupervised, while those for present-day Faroese individuals were estimated semi-supervised using ancient genomes as references. The five colors represent different ancestral sources: orange for West Europe, green for North Europe, blue for Steppe, purple for the Levant and East Mediterranean, and red for East Europe. The geographical distribution (bottom-right) highlights historical samples (250 years BP) in red, this study’s samples in black, and an 800-year-old individual sample in blue.

We estimated admixture timing for modern (n = 40) and ancient (n = 11) Faroese individuals using *DATES* with North and West Europe proxy references.^65^ The ancient Faroese had an average admixture estimate of 94.665 ± 58.658 generations prior to the dated age of the samples (∼980 BCE; 2,681 BCE - 720 CE, assuming 29-year generations) (Fig. S8A), while present-day Faroese showed more recent admixture at 72.567 ± 15.290 generations in the past (∼137 BCE; 581 BCE - 306 CE) (Fig. S8B). The high standard errors and inconsistency between estimates may reflect confounding due to drift and bottlenecks in the target population (as noted by Narasimhan, Patterson et al.)^65^ or low differentiation in linkage patterns between the source populations. Additionally, *DATES* assumes a single admixture pulse, but additional waves, particularly in the present-day Faroese, could shift estimates toward the present. In both cases, estimated admixture timing predates Faroese settlement, which likely began in the 9th century CE,^1–3^ but possibly as early as the 4th-6th centuries CE.^4^

## Discussion

Here we have presented the first whole-genome sequence data from 40 minimally related individuals from across the Faroe Islands, a North Atlantic founder population. The Faroese have a high prevalence of several diseases in comparison to other European or global populations, several of which have been of particular interest in epidemiological studies of the region (e.g. inflammatory bowel disease, type 2 diabetes, multiple sclerosis).^9–18^ Additionally, observational evidence from the FarGen project recruitment data suggest a higher prevalence of ankylosing spondylitis, although follow-up epidemiological studies are required. We investigated the frequency of HLA-B27 in the Faroe Islands, an allele which has been previously associated with ankylosing spondylitis (AS). We found that despite the observed high prevalence of AS in the Faroe Islands there was no evidence of increased HLA-B27 allele frequency compared to other European populations. However, a stop gain variant in *SERPINB10* was among those enriched in the Faroese cohort, and it may contribute to increased AS risk (rs138084090, AF = 2.5%). Rare variant analysis of this gene in the UK Biobank found that an independent stop gain variant in the same gene is nominally associated with increased AS risk (rs145346731, p = 2.5e-4), which was the most significant phenotypic association for the gene.^66^ *SERPINB10* is most highly expressed in neutrophilic metamyelocytes,^67^ a cell type with disease relevance to AS.^68^ *CCDC168*, another gene with a Faroese enriched stop gain variant had highly significant rare variant associations in the UK Biobank with corneal hysteresis and intraocular pressure (rs1361247423, p = 1.79e-57, p = 5.21e-13, respectively). Finally, a Faroese enriched stop gain variant in *AGL* identified in this study (rs113994128) has been previously reported as causing glycogen storage disease type IIIA in the Faroe Islands, which is estimated to have the highest prevalence of the disease world-wide.^69^ These findings emphasize the importance of further research into the role that unique genetic variation in the Faroe Islands may play in the incidence of diseases that, while common in the Faroes, have a global burden as well.

We found elevated amounts of the genome contained in long runs of homozygosity (ROHs) compared to other European reference cohorts, including another founder population from Finland. The higher proportion of the Faroese genome contained in these long ROHs suggests a stronger or more recent bottleneck in the Faroese population history. With the second phase of the FarGen study, a larger sample size will facilitate investigation into the timing and severity of the bottleneck(s). Additionally, founder populations such as the Finnish, have been a common focus for studies of founder events, genetic isolation, and the effects of haplotype sharing on aspects of human health and disease.^70–73^ As such, the Faroese population may serve as another useful global reference for studying the influence of demographic history on genetic variation and trait architecture. Long ROHs, in particular, can be enriched for deleterious variation or be of interest in understanding the genetic architecture of health-related traits in human populations.^39,74^ Studying these long ROHs may be relevant for future studies of health outcomes or other traits of interest in the Faroese population.

We also detected several regions under recent positive selection in the population. It is likely that iHS measures selection older than the settlement of the Faroe Islands. Indeed, we found that many of the top selection signals were shared between the Faroese and British cohorts, which is unsurprising given the recent divergence between these two populations and likely reflecting selection that began prior to this divergence. We did find differences in the strength of these signals; for example, there is more diversity on the Faroese *LCT*/*MCM6* lactase persistence haplotypes. The lactase persistence allele rs4988235 (chr2_135851076_G_A) has been inferred to be under strong selection at least until the medieval period in northwestern European groups.^75,76^ Patterson et al. 2021 found that the rate of increase in allele frequency may have slowed in recent periods; however, this does not exclude the possibility of continued or fluctuating selective pressure as this is consistent with the expected sigmoidal trajectory for an allele under ongoing selection.^77^ While the increased diversity on the Faroese lactase persistence haplotypes may be simply explained by population-specific drift, this result could also indicate relaxed selection for lactase persistence alleles after settlement of the Faroe Islands, possibly due to changes in dietary habits in the new environment. The traditional diet of the Faroe Islands consisted of a higher reliance on animal and marine fats such as sheep tallow, whale blubber, and liver from codfishes, while dairy products such as milk and cheese, and particularly that from cattle, were more limited in availability.^78^ The selected variant at rs4988235 is at 74% frequency in the modern Faroese cohort, and based on the ancient genomes for which we have available data, we can attest that the allele was present at in the Faroe Islands at high frequencies already in the 17th/18th centuries (∼82%), and imputation further suggests the haplotype containing the allele to at least have been present in the islands 800 years ago. We caveat that the sample size of the historical Faroese (11 individuals) is small and coverage of ancient samples is low, leading to potential errors in imputation. In the absence of selection or drift, we can calculate the expected frequency of an allele in an admixed population as the linear combination of allele frequencies and average ancestry contributions from the sources. Based on the frequency of the rs4988235 variant in proxy sources from the ancient panel, the expected allele frequency in the ancestral population at the time of admixture is approximately 47%. The difference in observed and expected allele frequencies may be due to drift, demography, changes in selection pressure, or a combination of these and other factors. We note limitations in this calculation as the proxy samples may not be good representatives for the true sources at the time of admixture, and there may have been multiple admixture events rather than a single pulse.

We detected selection targets that were specific to the Faroese population using the XPEHH statistic with the British cohort as the comparison population. As XPEHH has the best power to identify alleles that are fixed or approaching fixation in one population but not others, it is unlikely to detect older selection events or incomplete sweeps from shared ancestral populations. One top selection signal is in *POLQ*, which plays a role in DNA repair and various cancers. Without collecting relevant phenotypes or environmental factors, it is difficult to hypothesize what selection pressure may be driving the strong signal in *POLQ*, so this will be an important area of follow-up for future studies. In another top signal, we find *SLC10A1* which plays a role in fat and vitamin D absorption. Positive selection related to differences in dietary fat intake has been hypothesized in many human populations, such as the Inuit population in Greenland.^79,80^ Also situated in a far northern latitude, the Faroese diet is similar to that of the Inuit population, relying on animal and marine fats.^78,81^ The relationship between *SLC10A1* and vitamin D levels may also be relevant, as the northern latitudes of the Faroe Islands and minimal UV exposure can lead to vitamin D deficiency, which has been hypothesized to be a strong selection pressure in populations in extreme latitudes.^82,83^

Although we hypothesize that these results suggest possible adaptations to environmental pressures of diet or UV exposure in northern latitudes, we cannot draw definitive conclusions based on this current study. It is certainly possible that variants that have risen to high frequencies due to past or ongoing positive selection now play a role in health outcomes in modern populations. For example, the gene *TBC1D4*, which was shown to be under positive selection in the Greenlandic Inuit population likely due to a historical diet low in carbohydrates, has been associated with type 2 diabetes and insulin resistance in the same population.^81,84^ The prevalences of some diseases enriched in the Faroese population may be related to genomic regions under positive selection. For example, the results of several studies have suggested a role of vitamin D deficiency in the development of multiple sclerosis.^85–87^ Future studies could involve collection of relevant phenotypes and focus on characterizing selective pressures and fine-mapping targets of selection as have been done in studies that more thoroughly characterized selection signals related to dietary adaptation and UV exposure and their functional consequences in other northern latitude populations.^80–82,84^

We have inferred ancestry tracts in the present day Faroese genomes that were inherited from ancient populations throughout Europe. We found that present-day Faroese individuals have similar relative ancestry contributions from past “North” and “West Europe” Iron Age populations. The most ancient genome available from the Faroe Islands matches ancestry patterns found in Iron Age West Europe. Admixture could have occurred either via a mixture of the original “West Europe” ancestry with individuals of predominantly “North Europe” ancestry, or a by replacement with individuals that were already of mixed ancestry at the time of arrival in the islands (the latter are not uncommon in Viking Age mainland Europe). Our analysis also suggests a bottleneck or a more progressive differentiation process in the islands relative to the mainland, which may postdate the most ancient Faroese genome currently available (approximately 800 years old). The most ancient Faroese sample from Margaryan *et al.* - composed almost entirely of “West Europe” ancestry - is a male individual found in a chapel-site in Suðuroy. Consistent with this, a local legend suggests this site may have been occupied by Irish monks.^88^ We note that it is difficult to draw conclusions based on a single individual. It is possible that this particular individual moved to the Faroe Islands within their lifetime. More ancient and present-day samples from the islands could shed further light on the history of the Faroese population.

The average admixture timing between “North Europe” and “West Europe” sources (as estimated by *DATES*) pre-date the settlement of the Faroe Islands (137 BCE - 980 BCE). This is consistent with the low variance in ancestry proportion within the Faroese individuals (both historical and modern), indicating enough time for recombination to break up long ancestry tracts and for global ancestry proportions to reach an equilibrium in the population. That is, these ancestry patterns, combined with the *DATES* estimates, suggest that the present-day Faroese population is most likely descended from already admixed founders who arrived on the islands. Importantly, estimates of admixture timing had high statistical noise, possibly due to several confounding factors including drift, demography, and low differentiation between sources. In particular, it is unclear how the bottleneck history of the Faroese population may affect the performance of *DATES*. In future studies, it will be informative to estimate and simulate the bottleneck size in the Faroese population, and then test the performance of *DATES* on those simulations to confirm whether bottleneck history has affected the empirical estimates of admixture timing. Additionally, it will be important to model single-pulse versus multiple pulses of admixture to determine whether this has resulted in the different estimates for admixture timing in modern and ancient Faroese.

This study focused on population genomic analyses such as selection scans, population structure, kinship, and ancestry. Given the unique settlement history and genetic architecture of the Faroe Islands, future studies which combine genomic data with relevant phenotype data could provide useful insight into the underlying genetic mechanisms of those traits. In particular, larger-scale genomic studies in the Faroese could investigate genetic risk factors which contribute to the high prevalence of autoimmune and metabolic disease on the islands. This is a focus of the second phase of the FarGen study, which is currently ongoing.

## Methods

### Sample Selection and Cryptic Relatedness

#### FarGen cohort

The participants in this study voluntarily enrolled in the FarGen project (The Faroe Genome Project: https://www.fargen.fo/en/home/). The 1,541 subjects are extensively described in Apol *et al.* 2022.^21^ Participant inclusion criteria for the FarGen project are that participants must live in the Faroe Islands or be of Faroese descent. Apol *et al.* report that 96.4% of the participants have between one and four Faroese grandparents. The cohort has a mixed health status composition, with 75% of the participants self-reporting that they have a confirmed diagnosis. Apol *et al.* found that the cohort is somewhat biased in terms of geographical representation, with the capital region being substantially over-represented.

#### Reconstruction of genealogy

The Multi-Generation Register at the Faroese Health Authority describes the ancestry of inhabitants of the Faroe Islands (http://fargen.fo/research/multi-generation-registry). The lineages can be traced back to approximately 1650 C.E. The register records birth date, parent identities, parents’ residence at the time of birth, and more. The Legacy Family Tree (https://legacyfamilytree.com) genealogy software is used to manage the digitized records. We reconstructed a genealogical tree of all the individuals in the FarGen cohort by looking up each individual, and recursively looking up their parents until there are no more ancestors. After reconstructing the genealogy of each individual two generations in the past, we discarded any who had fewer than 6 direct ancestors recorded in the Multi-Generation Registry (2 parents and 4 grandparents). We note that although an individual is recorded in the register, there is no guarantee they were born in the Faroe Islands.

#### Geographical stratification through dialect

We defined six geographical regions of the Faroe Islands as annotated in Fig. 1A: Norðoyggjar; Eysturoy and Norðstreymoy; Suðurstreymoy; Vágar and Mykines; Sandoy, Skúvoy, and Stóra Dímun; Suðuroy, and placed individuals within these regions based on birth place. The boundaries of the regions were defined using isoglosses (i.e. boundaries where we see changes in dialect) as described in Þráinsson 2012.^89^ For example, the isogloss for "á", which may be pronounced either as [a:] or [ɔa], separates Norðoyggjar in the north from the rest of the islands.

#### Calculating pairwise kinship coefficients

In order to avoid sequencing highly related samples, we used the large constructed genealogy to account for cryptic relatedness. We calculated pairwise kinship coefficient between every individual using *kinship2*.^90^ This method assumes that the founders (individuals in the pedigree without recorded parents) are unrelated to other founders and each individual founder’s parents were not related to each other, which may not always be the case in this population.

#### Sample selection using graph theory

We constructed a relationship graph with nodes representing individuals, and connected two nodes by an edge if their kinship coefficient is above a given threshold as described below. Ideally, we would remove nodes such that all edges are removed, while keeping as many nodes as possible, referred to as the *maximum independent set* problem. Obtaining an exact solution to the maximum independent set is an NP-hard problem (exponential time complexity), making it infeasible for our applications. Instead, we obtained an approximate maximum independent set using an algorithm described in Boppana *et al.* 1990.^91^ We performed relatedness pruning with a threshold of 2^−6^ on 1,294 FarGen individuals who were not missing a birth region, resulting in 332 individuals. This was the minimum threshold for which we could have enough sampling from each region. From these 332 individuals, we sampled 5 to 8 from each region, yielding 40 individuals for whole genome sequencing and subsequent analyses.

### Bioinformatics

#### Sequencing and quality control

DNA samples from the 40 selected individuals were sequenced at FarGen (using TruSeq PCR-free libraries on Illumina NextSeq 500 instruments) to an average depth of 19.2x, ranging from 9.9x to 31.5x per sample. Sequencing QC was investigated with FASTQC, Picard (including CollectWgsMetrics for coverage) and VerifyBamID for contamination.

#### Variant calling and imputation

Reads were processed with Variant Bio’s in-house processing pipeline based on the GATK Best Practices (CCDG functional equivalence version).^92^ Joint genotyping was performed with GATK (version 4.2.0.0) including 714 genomes from the 1000 Genomes Project (503 Europeans, 103 Han Chinese, 108 Yoruba) and followed by VQSR (--truth-sensitivity-filter-level to 99.8 for SNPs and 99.0 for indels). Only PASS variants in GIAB high-confidence regions (∼80% of GRCh38) were retained.^93^ Genotypes with GQ<=20 were filtered (set to missing) and imputation was performed within the full cohort of 754 genomes using Beagle v5.1. Variants enriched in the Faroese cohort and with predicted functional impact (Table S2) were additionally hard-filtered with VQSLOD>20.

#### HLA typing

We ran HLA*LA (v1.0.3) on the mapped reads to determine HLA types for the 40 Faroese individuals as well as for the GBR, CEU, and FIN reference population individuals from 1000 Genomes.^29^ Benchmarking the method on 1000 Genomes Project data, where HLA types are known, we estimate overall HLA-B typing accuracy at 93.4%, 88.8% and 89.9% for the GBR (N=91), CEU (N=99), and FIN (N=99) reference populations. Accuracy of B27 detection specifically is 100% (N=10), 83.3% (N=6) and 100% (N=15) based on these three reference cohorts, respectively, with one B*27:05 allele mis-identified as B*27:26 in CEU. Table S3 contains counts, mean quality scores, and frequencies of all HLA-B alleles detected among the 40 Faroese as well as the GBR, CEU, and FIN reference populations.

### Population Genetics

#### Individual and variant-level filtering

Beginning with a total of 21,837,577 variants and 754 individuals, we applied various individual-level and variant-level quality control filters for downstream analyses. We filtered 8 individuals with mismatched sex based on genetics and reported information. We additionally filtered 5 individuals that were, for any of PCs 1-10, further than 7 standard deviations from the mean. We then filtered 481,403 variants that were not in GIAB high confidence regions, had MSQ rank sum not equal to zero or failed gnomAD v3 QC.^94^ We removed 3,627 variants with a minor allele count of less than 1 after individual-level filters were applied. We removed 695,947 variants that were not autosomal. This final dataset included 20,656,600 variants and 741 individuals.

#### Selection scans

iHS and XPEHH were calculated using the *hapbin* software (https://github.com/evotools/hapbin) with option –max-extend 1000000 and --minmaf 0.05.^41^ All other options were set to default. To compute P-values, we used the method by Fariello et al. (2013), exploiting the fact that detectable regions under strong selection affect a small portion of the genome.^95^ For both statistics, values were standardized genome-wide in 2% allele frequency bins, as allele frequency is correlated with allele age and therefore haplotype length.^40,43^ We first computed outlier-robust mean and standard deviation with the rlm() function from the MASS package in *R*, to reduce the influence of outliers.^95,96^ The standardized values of these summary statistics represent z-scores. We calculated two tailed p-values using these z-scores, giving the probability that we observe these data by chance compared to null expectations for the standard normal distribution. Q-Q plots and histograms of p-values for each statistic are provided in the Supplementary Materials (Fig. S9). For each summary statistic distribution, we also calculated the p-value cutoffs that correspond to a False Discovery Rate (FDR) of less than 0.01 and 0.001, using the q-value *R* package (https://github.com/StoreyLab/qvalue).^42^ We additionally include the empirical standardized values for each statistic (Fig. S10).

EHH decay plots and haplotype furcations for the *LCT* locus were calculated using the *rehh R* package (https://cran.r-project.org/web/packages/rehh/index.html)^47^ and visualized the haplotype structure of the genomic region chr2:135701076-136009184 using *haplostrips* and plot option -S 3 (https://bitbucket.org/dmarnetto/haplostrips).^48^

#### Kinship and runs of homozygosity

The kinship matrix in the WGS cohort was calculated using the *popkin* software (https://github.com/StoreyLab/popkin).^36–38^ We restricted the analyses to biallelic SNPs with a minor allele frequency of at least 0.01 in at least one subpopulation (i.e. YRI, CHB, FIN, CEU, GBR, TSI, IBS, or FARO), resulting in a dataset of 15,206,409 variants. When calculating the kinship matrix for the Faroese WGS cohort only, we used the *rescale_kinship()* function, which will change the most recent common ancestor and give different absolute values, but the overall relationship structure in the subpopulation remains the same. Using the same data set, we calculated runs of homozygosity (ROHs) for each individual using *bcftools/RoH*.^97^ “Short” ROHs were classified as ROH less than or equal to 300 kb, “medium” as greater than 300kb and less than or equal to 1 Mb, and “long” as greater than 1 Mb.

#### Fine-scale structure estimation using ancient genomes

A panel of 616 imputed ancient genomes from Allentoft *et al.* 2024 (downloaded from https://doi.org/10.17894/ucph.d71a6a5a-8107-4fd9-9440-bdafdfe81455), representing individuals from several European regions (southern Europe, western Europe, northern Europe, eastern Europe, and central Europe) was used for analyses.^63^ Only samples that were estimated to be no older than 3,000 years old were used. Out of these ancient samples, 11 were excavated in the Faroe Islands, 10 of them are historical samples dated to approximately 250 years old, and 1 of them is dated to be approximately 800 years old.^64^ The sample location, approximate age, and sources for these samples are listed in Table S5. To consolidate the two panels, we first performed a liftover of the ancient genome VCF files to the GRCh38 reference genome. Following this, we applied quality filters to the dataset (bi-allelic sites MAF > 0.05 and imputation INFO >= 0.5).

We used HaploNet - a neural network-based method for performing window-based haplotype clustering across the genome - for fine-scale population structure inference on the combined panel. HaploNet uses a hidden Markov model to find an optimal window-based local ancestry “painting” across a genome, given estimated haplotype cluster likelihoods, haplotype cluster frequencies, and global ancestry proportions.^98^ The Faroese panel’s haplotype frequencies are very homogeneous and highly differentiated from mainland Europeans. For this reason, under an initial round of unsupervised ancestry estimation, we found that the Faroese individuals captured a major component at first split (K=2). We therefore implemented and utilized a semi-supervised ancestry estimation feature in HaploNet^20^. We performed haplotype clustering in non-overlapping windows of 512 SNPs, and we used the resulting haplotype cluster likelihoods to perform principal component analysis (PCA) and estimate both global and local ancestry in the Faroese individuals.

We performed global ancestry estimation in HaploNet using its EM algorithm to find the maximum likelihood estimates using only ancient European individuals (excluding the Faroese individuals).^20^ We used the EM algorithm a second time to estimate the ancestry proportions (Q matrix) in the Faroese individuals. The estimated haplotype cluster frequencies (F matrix) were kept fixed, which means that the semi-supervised approach can be seen as modeling the Faroese individuals using inferred haplotype clusters from the ancient European individuals.

We estimated the average timing of admixture in modern and ancient Faroese individuals using the *DATES* software.^65^ We selected reference individuals from the ancient panel who were maximized for North Europe (n=64) and West Europe (n=41) ancestry, respectively. We obtained separate estimates for the admixture timing in the 11 historical Faroese individuals dated to approximately the 17th century and the 40 modern individuals from the FarGen project. We used the following recommended default options for optimal performance: binsize = 0.001, maxdis = 1.0, jacknife = YES, qbin = 10, runfit = YES, afffit = YES, lovalfit = 0.45.

## Data Availability

Variant-level summary statistics and genome-wide selection scan results for iHS and XPEHH are available for research via AWS S3 (s3://<u>public.us-prod.variantbio.com/FaroeIslands_SelectionScans/</u>)l. Genetic and meta data from this study is stored at the Faroese Health Authority. Access to individual-level data is available for research upon participants’ re-consent. Researchers will be granted access to de-identified genetic data and metadata, provided that the project protocol has been approved by the Faroese Scientific Ethical Committee and a template material/data transfer agreement has been signed with the Faroese Health Authority in compliance with GDPR (see Gregersen *et al.*, 2021). Requests should be made to Noomi O. Gregersen.

## Acknowledgements

We thank the participants of the FarGen project. FarGen is supported by the Government of the Faroe Islands. F.R. is supported by a Novo Nordisk Fonden Data Science Ascending Investigator Award (NNF22OC0076816) and by the European Research Council (ERC) under the European Union’s Horizon Europe programme (grant agreements 101077592 and 951385). We also thank Victor Lee with assistance while working with ancient genomic data.

## Author Contributions

S.E.C., K.A.W., F.R. and N.O.G. designed the study. N.O.G. and K.D.A. secured ethical permissions, facilitated the inclusion of participants, and oversaw the compliance with ethical standards and protocols. G.A. prepared the data for the genealogy analysis. Ó.M. performed relatedness pruning and sample selection for WGS. L.N.L prepared samples for WGS and performed the WGS analyses. A.E. processed sequencing data, performed imputation, and carried out quality control. I.H. conducted kinship, ROH, and selection scan analyses. A.R.M. carried out ancient admixture and local ancestry analysis. J.M. added functionalities to Haplonet software for semi-supervised admixture and fine-structure analysis. S.E.C., N.M., and F.R. supervised analyses. I.H., S.E.C., and A.R.M. interpreted results and wrote the manuscript. All authors reviewed and contributed to the writing of this manuscript.

## Materials & Correspondence

Correspondence to N.O.G., S.E.C., and F.R..

## References

1. Johnston, G. The Faroe Islanders Saga. (Oberon Books, Canada, 1975).

2. Jorgensen, T. H. et al. The origin of the isolated population of the Faroe Islands investigated using Y chromosomal markers. Hum. Genet. 115, 19–28 (2004).

3. Young, G. V. C. *From the Vikings to the Reformation: A Chronicle of the Faroe Islands Up to* 1538. (Nám, 1979).

4. Church, M. J. et al. The Vikings were not the first colonizers of the Faroe Islands. Quat. Sci. Rev. 77, 228–232 (2013).

5. Als, T. D. et al. Highly discrepant proportions of female and male Scandinavian and British Isles ancestry within the isolated population of the Faroe Islands. Eur. J. Hum. Genet. 14, 497–504 (2006).

6. West, J. F. Faroe: The Emergence of a Nation. (C. Hurst, London, 1972).

7. Jorgensen, T. H. et al. Linkage disequilibrium and demographic history of the isolated population of the Faroe Islands. Eur. J. Hum. Genet. 10, 381–387 (2002).

8. Gislason, H. SNP heterozygosity, relatedness and inbreeding of whole genomes from the isolated population of the Faroe Islands. BMC Genomics 24, 707 (2023).

9. Leblond, C. S. et al. Both rare and common genetic variants contribute to autism in the Faroe Islands. Npj Genomic Med. 4, 1–10 (2019).

10. Rasmussen, J. et al. Carnitine levels in 26,462 individuals from the nationwide screening program for primary carnitine deficiency in the Faroe Islands. J. Inherit. Metab. Dis. 37, 215–222 (2014).

11. Passa, P. Diabetes trends in Europe. Diabetes Metab. Res. Rev. 18 **Suppl 3**, S3–8 (2002).

12. Veyhe, A. S. et al. Prevalence of type 2 diabetes and prediabetes in the Faroe Islands. Diabetes Res. Clin. Pract. 140, 162–173 (2018).

13. Burisch, J. et al. East–West gradient in the incidence of inflammatory bowel disease in Europe: the ECCO-EpiCom inception cohort. Gut 63, 588–597 (2014).

14. Dean, L. E. et al. Global prevalence of ankylosing spondylitis. Rheumatology 53, 650–657 (2014).

15. Schwartz, M., Sørensen, N., Brandt, N. J., Høgdall, E. & Holm, T. High incidence of cystic fibrosis on The Faroe Islands: a molecular and genealogical study. Hum. Genet. 95, 703–706 (1995).

16. Joensen, P. Multiple sclerosis: variation of incidence of onset over time in the Faroe Islands. Mult. Scler. J. 17, 241–244 (2011).

17. Hammer, T., Nielsen, K. R., Munkholm, P., Burisch, J. & Lynge, E. The Faroese IBD Study: Incidence of Inflammatory Bowel Diseases Across 54 Years of Population-based Data. J. Crohns Colitis 10, 934–942 (2016).

18. Gregersen, N. O. et al. Whole-exome sequencing implicates DGKH as a risk gene for panic disorder in the Faroese population. Am. J. Med. Genet. B Neuropsychiatr. Genet. 171, 1013–1022 (2016).

19. Gregersen, N. O., Apol, K. D., Weihe, P., Steig, B. Á & Andorsdóttir, G. FarGen: Bioresource From the Faroe Genome Project. 8, 1 (2021).

20. Meisner, J. & Albrechtsen, A. Haplotype and population structure inference using neural networks in whole-genome sequencing data. Genome Res. gr.276813.122 (2022) doi:10.1101/gr.276813.122.

21. Apol, K. D. et al. FarGen – participants in the genetic research infrastructure of the Faroe Islands. Scand. J. Public Health 50, 980–987 (2022).

22. Emde, A.-K. et al. Mid-pass whole genome sequencing enables biomedical genetic studies of diverse populations. BMC Genomics 22, 666 (2021).

23. Auton, A. et al. A global reference for human genetic variation. Nature 526, 68–74 (2015).

24. Goyette, P. et al. High-density mapping of the MHC identifies a shared role for HLA-DRB1*01:03 in inflammatory bowel diseases and heterozygous advantage in ulcerative colitis. Nat. Genet. 47, 172–179 (2015).

25. Braun, J. & Sieper, J. Fifty years after the discovery of the association of HLA B27 with ankylosing spondylitis. RMD Open 9, e003102 (2023).

26. Parameswaran, P. & Lucke, M. HLA-B27 Syndromes. in StatPearls (StatPearls Publishing, Treasure Island (FL), 2024).

27. Bowness, P. HLA-B27. Annu. Rev. Immunol. 33, 29–48 (2015).

28. Hanson, A. & Brown, M. A. Genetics and the causes of ankylosing spondylitis. Rheum. Dis. Clin. North Am. 43, 401–414 (2017).

29. Dilthey, A. T. et al. HLA*LA—HLA typing from linearly projected graph alignments. Bioinformatics 35, 4394–4396 (2019).

30. Gonzalez-Galarza, F. F. et al. Allele frequency net database (AFND) 2020 update: gold-standard data classification, open access genotype data and new query tools. Nucleic Acids Res. 48, D783–D788 (2020).

31. Hurley, C. K. et al. Common, intermediate and well-documented HLA alleles in world populations: CIWD version 3.0.0. HLA 95, 516–531 (2020).

32. Robinson, J. et al. IPD-IMGT/HLA Database. Nucleic Acids Res. 48, D948–D955 (2020).

33. Dillon, C. F. & Hirsch, R. The United States National Health and Nutrition Examination Survey and the epidemiology of ankylosing spondylitis. Am. J. Med. Sci. 341, 281–283 (2011).

34. Cortes, A. et al. Major histocompatibility complex associations of ankylosing spondylitis are complex and involve further epistasis with ERAP1. Nat. Commun. 6, 7146 (2015).

35. Brown, M. A. et al. HLA class I associations of ankylosing spondylitis in the white population in the United Kingdom. Ann. Rheum. Dis. 55, 268–270 (1996).

36. Ochoa, A. & Storey, J. D. Estimating FST and kinship for arbitrary population structures. PLOS Genet. 17, e1009241 (2021).

37. Ochoa, A. & Storey, J. D. New kinship and FST estimates reveal higher levels of differentiation in the global human population. 653279 Preprint at 10.1101/653279 (2019).

38. Ochoa, A. & Storey, J. D. FST and kinship for arbitrary population structures I: Generalized definitions. 083915 Preprint at 10.1101/083915 (2019).

39. Ceballos, F. C., Joshi, P. K., Clark, D. W., Ramsay, M. & Wilson, J. F. Runs of homozygosity: windows into population history and trait architecture. Nat. Rev. Genet. 19, 220–234 (2018).

40. Voight, B. F., Kudaravalli, S., Wen, X. & Pritchard, J. K. A Map of Recent Positive Selection in the Human Genome. PLOS Biol. 4, e72 (2006).

41. Maclean, C. A., Chue Hong, N. P. & Prendergast, J. G. D. hapbin: An Efficient Program for Performing Haplotype-Based Scans for Positive Selection in Large Genomic Datasets. Mol. Biol. Evol. 32, 3027–3029 (2015).

42. Storey, J. D. & Tibshirani, R. Statistical significance for genomewide studies. Proc. Natl. Acad. Sci. 100, 9440–9445 (2003).

43. Sabeti, P. C. et al. Genome-wide detection and characterization of positive selection in human populations. Nature 449, 913–918 (2007).

44. Fan, S., Hansen, M. E. B., Lo, Y. & Tishkoff, S. A. Going global by adapting local: A review of recent human adaptation. Science 354, 54–59 (2016).

45. Williamson, S. H. et al. Localizing Recent Adaptive Evolution in the Human Genome. PLoS Genet. 3, e90 (2007).

46. Sabeti, P. C. et al. Detecting recent positive selection in the human genome from haplotype structure. Nature 419, 832–837 (2002).

47. Gautier, M. & Vitalis, R. rehh: an R package to detect footprints of selection in genome-wide SNP data from haplotype structure. Bioinformatics 28, 1176–1177 (2012).

48. Marnetto, D. & Huerta-Sánchez, E. Haplostrips: revealing population structure through haplotype visualization. Methods Ecol. Evol. 8, 1389–1392 (2017).

49. Floerl, S. et al. Functional and Pharmacological Comparison of Human and Mouse Na+/Taurocholate Cotransporting Polypeptide (NTCP). SLAS Discov. Adv. Life Sci. R D 26, 1055–1064 (2021).

50. Vaz, F. M. et al. Sodium taurocholate cotransporting polypeptide (SLC10A1) deficiency: conjugated hypercholanemia without a clear clinical phenotype. Hepatol. Baltim. Md 61, 260–267 (2015).

51. Ho, R. H., Leake, B. F., Roberts, R. L., Lee, W. & Kim, R. B. Ethnicity-dependent polymorphism in Na+-taurocholate cotransporting polypeptide (SLC10A1) reveals a domain critical for bile acid substrate recognition. J. Biol. Chem. 279, 7213–7222 (2004).

52. Hagenbuch, B. & Meier, P. J. Molecular cloning, chromosomal localization, and functional characterization of a human liver Na+/bile acid cotransporter. J. Clin. Invest. 93, 1326–1331 (1994).

53. Deng, M. et al. Clinical and molecular study of a pediatric patient with sodium taurocholate cotransporting polypeptide deficiency. Exp. Ther. Med. 12, 3294–3300 (2016).

54. Liu, R. et al. Homozygous p.Ser267Phe in SLC10A1 is associated with a new type of hypercholanemia and implications for personalized medicine. Sci. Rep. 7, 9214 (2017).

55. Ceccaldi, R. et al. Homologous-recombination-deficient tumours are dependent on Polθ-mediated repair. Nature 518, 258–262 (2015).

56. Yoon, J.-H., Roy Choudhury, J., Park, J., Prakash, S. & Prakash, L. A role for DNA polymerase θ in promoting replication through oxidative DNA lesion, thymine glycol, in human cells. J. Biol. Chem. 289, 13177–13185 (2014).

57. Arana, M. E., Seki, M., Wood, R. D., Rogozin, I. B. & Kunkel, T. A. Low-fidelity DNA synthesis by human DNA polymerase theta. Nucleic Acids Res. 36, 3847–3856 (2008).

58. Seki, M., Marini, F. & Wood, R. D. POLQ (Pol theta), a DNA polymerase and DNA-dependent ATPase in human cells. Nucleic Acids Res. 31, 6117–6126 (2003).

59. Wood, R. D. & Doublié, S. DNA polymerase θ (POLQ), double-strand break repair, and cancer. DNA Repair 44, 22–32 (2016).

60. Thomas, C. et al. Melanoma-derived DNA polymerase theta variants exhibit altered DNA polymerase activity. BioRxiv Prepr. Serv. Biol. 2023.11.14.566933 (2023) doi:10.1101/2023.11.14.566933.

61. Yoon, J.-H. et al. Error-Prone Replication through UV Lesions by DNA Polymerase θ Protects against Skin Cancers. Cell 176, 1295–1309.e15 (2019).

62. Pan, Q. et al. Knockdown of POLQ interferes the development and progression of hepatocellular carcinoma through regulating cell proliferation, apoptosis and migration. Cancer Cell Int. 21, 482 (2021).

63. Allentoft, M. E. et al. Population genomics of post-glacial western Eurasia. Nature 625, 301–311 (2024).

64. Margaryan, A. et al. Population genomics of the Viking world. Nature 585, 390–396 (2020).

65. Narasimhan, V. M. et al. The formation of human populations in South and Central Asia. Science 365, eaat7487 (2019).

66. Karczewski, K. J. et al. Systematic single-variant and gene-based association testing of thousands of phenotypes in 394,841 UK Biobank exomes. Cell Genomics 2, (2022).

67. George, N. et al. Expression Atlas update: insights from sequencing data at both bulk and single cell level. Nucleic Acids Res. 52, D107–D114 (2024).

68. Coletto, L. A. et al. The Role of Neutrophils in Spondyloarthritis: A Journey across the Spectrum of Disease Manifestations. Int. J. Mol. Sci. 24, 4108 (2023).

69. Santer, R. et al. Molecular genetic basis and prevalence of glycogen storage disease type IIIA in the Faroe Islands. Eur. J. Hum. Genet. 9, 388–391 (2001).

70. Norio, R. Finnish Disease Heritage II: population prehistory and genetic roots of Finns. Hum. Genet. 112, 457–469 (2003).

71. Kerminen, S. et al. Fine-Scale Genetic Structure in Finland. G3 GenesGenomesGenetics 7, 3459–3468 (2017).

72. Peltonen, L., Jalanko, A. & Varilo, T. Molecular Genetics the Finnish Disease Heritage. Hum. Mol. Genet. 8, 1913–1923 (1999).

73. Sabatti, C. et al. Genome-wide association analysis of metabolic traits in a birth cohort from a founder population. Nat. Genet. 41, 35–46 (2009).

74. Szpiech, Z. A. et al. Long runs of homozygosity are enriched for deleterious variation. Am. J. Hum. Genet. 93, 90–102 (2013).

75. Patterson, N. et al. Large-scale migration into Britain during the Middle to Late Bronze Age. Nature 601, 588–594 (2022).

76. Burger, J. et al. Low Prevalence of Lactase Persistence in Bronze Age Europe Indicates Ongoing Strong Selection over the Last 3,000 Years. Curr. Biol. 30, 4307–4315.e13 (2020).

77. Haldane, J. B. S. A Mathematical Theory of Natural and Artificial Selection, Part V: Selection and Mutation. Math. Proc. Camb. Philos. Soc. 23, 838–844 (1927).

78. Svanberg, I. The Importance of Animal and Marine Fat in the Faroese Cuisine: The Past, Present, and Future of Local Food Knowledge in an Island Society. Front. Sustain. Food Syst. 5, (2021).

79. Buckley, M. T. et al. Selection in Europeans on Fatty Acid Desaturases Associated with Dietary Changes. Mol. Biol. Evol. 34, 1307–1318 (2017).

80. Fumagalli, M. et al. Greenlandic Inuit show genetic signatures of diet and climate adaptation. Science 349, 1343–1347 (2015).

81. Andersen, M. K. & Hansen, T. Genetics of metabolic traits in Greenlanders: lessons from an isolated population. J. Intern. Med. 284, 464–477 (2018).

82. Hlusko, L. J. et al. Environmental selection during the last ice age on the mother-to-infant transmission of vitamin D and fatty acids through breast milk. Proc. Natl. Acad. Sci. 115, E4426–E4432 (2018).

83. Nielsen, R. et al. Tracing the peopling of the world through genomics. Nature 541, 302–310 (2017).

84. Moltke, I. et al. A common Greenlandic TBC1D4 variant confers muscle insulin resistance and type 2 diabetes. Nature 512, 190–193 (2014).

85. Salzer, J. et al. Vitamin D as a protective factor in multiple sclerosis. Neurology 79, 2140–2145 (2012).

86. Sintzel, M. B., Rametta, M. & Reder, A. T. Vitamin D and Multiple Sclerosis: A Comprehensive Review. Neurol. Ther. 7, 59–85 (2017).

87. Laursen, J. H. et al. Genetic and environmental determinants of 25-hydroxyvitamin D levels in multiple sclerosis. Mult. Scler. J. 21, 1414–1422 (2015).

88. Arge, S. V. Christianity, churches and medieval Kirkjubøur – contacts and influences in the Faroe Islands. in Medieval Archaeology in Scandinavia and Beyond: History, Trends and Tomorrow 235–256 (Aarhus University Press, 2015).

89. Þráinsson, H. Faroese: An Overview and Reference Grammar. (Føroya Fróđskaparfelag, 2004).

90. Sinnwell, J. P., Therneau, T. M. & Schaid, D. J. The kinship2 R package for pedigree data. Hum. Hered. 78, 91–93 (2014).

91. Boppana, R. & Halldórsson, M. M. Approximating maximum independent sets by excluding subgraphs. BIT Numer. Math. 32, 180–196 (1992).

92. Regier, A. A. et al. Functional equivalence of genome sequencing analysis pipelines enables harmonized variant calling across human genetics projects. Nat. Commun. 9, 4038 (2018).

93. Olson, N. D. et al. PrecisionFDA Truth Challenge V2: Calling variants from short and long reads in difficult-to-map regions. Cell Genomics 2, 100129 (2022).

94. Karczewski, K. J. et al. The mutational constraint spectrum quantified from variation in 141,456 humans. Nature 581, 434–443 (2020).

95. Fariello, M. I., Boitard, S., Naya, H., SanCristobal, M. & Servin, B. Detecting Signatures of Selection Through Haplotype Differentiation Among Hierarchically Structured Populations. Genetics 193, 929–941 (2013).

96. Venables, W. N. & Ripley, B. D. *Modern Applied Statistics with S*. (Springer, New York, NY, 2002).

97. Narasimhan, V. et al. BCFtools/RoH: a hidden Markov model approach for detecting autozygosity from next-generation sequencing data. Bioinformatics 32, 1749–1751 (2016).

98. Scheet, P. & Stephens, M. A Fast and Flexible Statistical Model for Large-Scale Population Genotype Data: Applications to Inferring Missing Genotypes and Haplotypic Phase. Am. J. Hum. Genet. 78, 629–644 (2006).

