## Supplementary Materials for "Faroese Whole Genomes Provide Insight into Ancestry and Recent Selection"

**Supplementary Tables** 2
**Supplementary Figures** 3

### Supplementary Tables

**Table S1. Whole genome sequencing quality control metrics.** Unique identifier (subject), sex, region, mean WGS depth (mean_depth), mean depth on X chromosome (mean_depth_chrX), mean depth on Y chromosome (mean_depth_chrY), median insert size (insert_median), mean absolute deviation of insert size (insert_mad), contamination estimate (contamination), and number of genotype calls with GQ > 20 (non_imputed_calls). The Faroes region labels are: VM = Vágar and Mykines; SR = Suðuroy; SM = Suðurstreymoy; SD = Sandoy, Skúvoy, Stóra Dímun; NG = Norðoyggjar; and EN = Eysturoy og Norðstreymoy.

**Table S2. Faroese putatively functional alleles.** Variants in the joint call set with CADD > 30 and at least two minor alleles observed in Faroese individuals, and no minor alleles observed in Finnish or Northern European reference individuals. Allele frequencies in Faroese individuals (af_FARO), allele frequencies in 1000 Genomes British (af_GBR), Central European (af_CEU), and Finnish (af_FIN), and allele frequencies in 1000G Europeans (af_1000G_EUR) and gnomAD (af_GNOMAD) references.

**Table S3. HLA-B Allele frequencies.** Counts of HLA-B alleles out of the total 80 haplotypes and mean quality scores in the Faroese WGS cohort, as well as HLA-B allele frequencies in three 1000 Genomes European cohorts (British [GBR], Central European [CEU], and Finnish [FIN]. HLA genotypes were determined using HLA*LA.

**Table S4. Genomic regions with strong signals of positive selection.** Genomic regions for the top 10 standardized iHS and XP-EHH values in the Faroese WGS cohort. We list the number of variants and all variant IDs in these regions which reach the p-value threshold corresponding to FDR < 0.01 (iHS: $p < 2.72 \times 10^{-6}$; XPEHH: $p < 2.35 \times 10^{-6}$). The regions are defined by the first and last variant position with a p-value below the cutoff for each particular statistic, as indicated in the “statistic” column. Protein-coding gene name(s), if any, with annotated variants reaching these thresholds are provided. We additionally provide the maximum absolute value of the given statistic (“max_statistic”), its corresponding p-value and q-value (i.e. the minimum FDR when that particular test is considered significant), and the allele frequencies in the Faroese and British cohorts for the variant(s) with the maximum value. The variant(s) with the maximum value for each region are highlighted in bold font.

**Table S5. Admixture and ancient ancestry samples.** Region, country, approximate age, publication source, and group label for ancient ancestry samples included in the ADMIXTURE and HaploNet analyses.

### Supplementary Figures

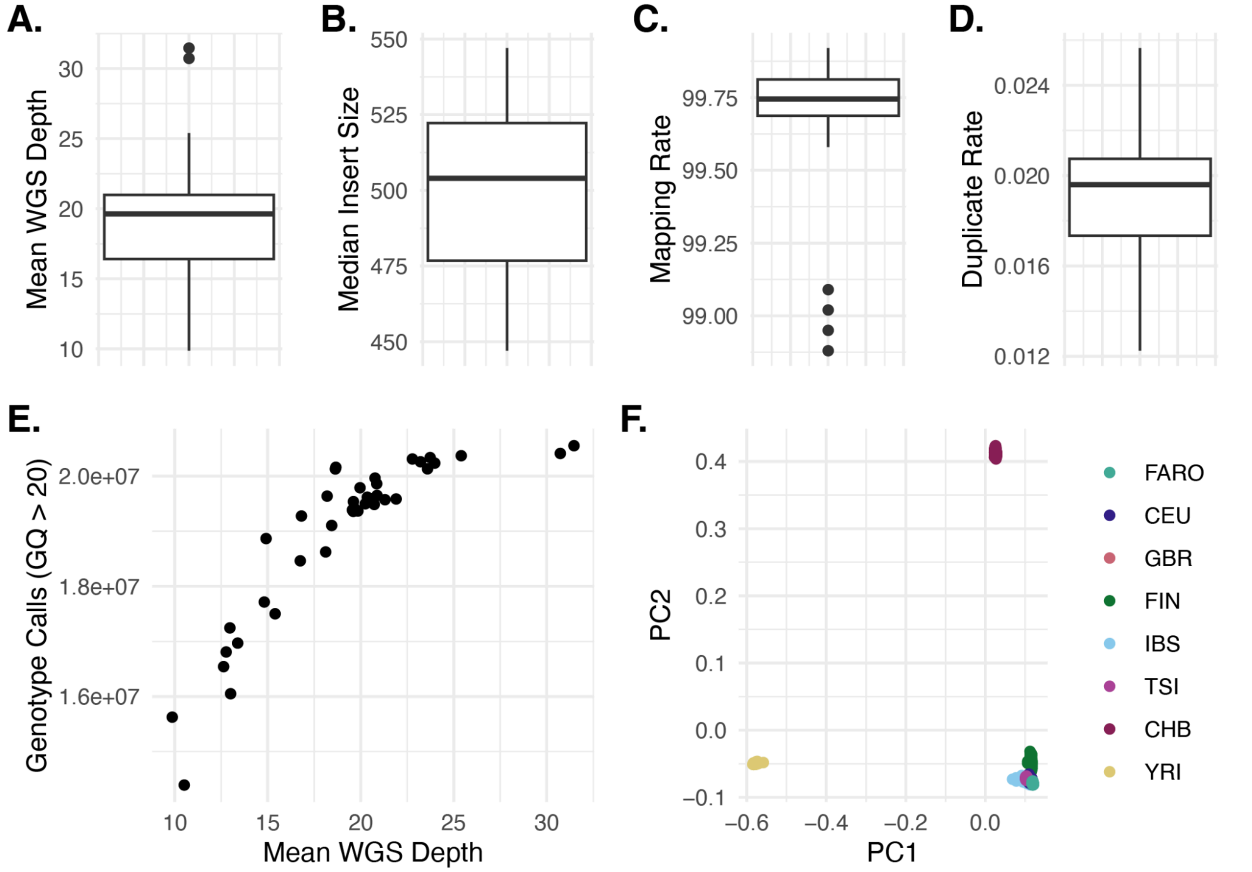


**Figure S1. Quality control of whole genome sequencing data. A-D)** Distributions of QC metrics across Faroese samples sequenced (mean depth, median insert size, mapping rate, duplicate rate). **E)** Mean WGS depth versus number of genotype calls with GQ > 20. Singletons and calls with GT <= 20 were discarded before imputation. **F)** Principal component analysis (PCA) of Faroese genomes jointly called with relevant 10000 Genomes reference data captures African ancestry in the first component and East Asian ancestry in the second component (FARO, Faroese, CEU, Central Europeans, GBR, British, FIN, Finnish IBS, Iberian, TSI, Tuscan, CHB, Han Chinese, YRI, Yoruban).

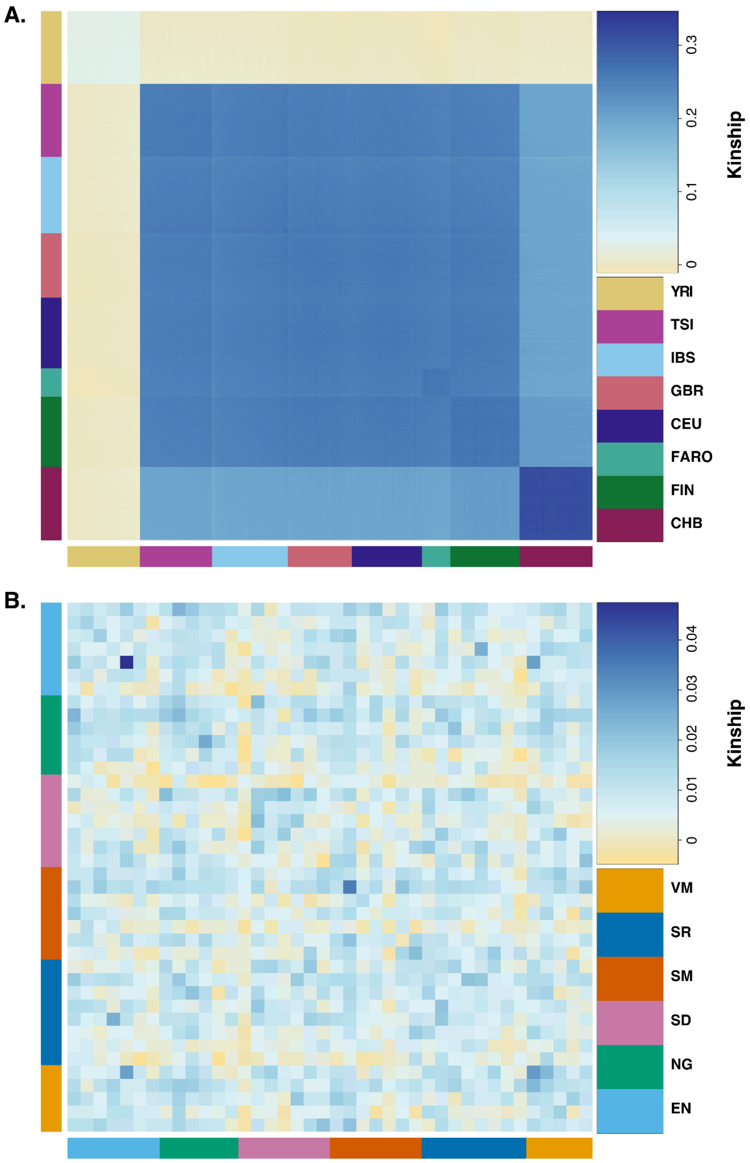


**Figure S2. Kinship matrices between modern populations. A)** Kinship estimated by *popkin*, including global reference populations from the 1000 Genomes and the Faroese WGS cohort (FARO). **B)** Kinship estimated by *popkin* for the Faroese WGS cohort only for better visualization. The matrix is rescaled after subsetting the individuals, so although the scales are different, the relative kinship within the cohort remains the same for the two plots. The matrices are symmetrical and ordered by population or region label as indicated by the colored bars along the rows and columns. The diagonal of each matrix is the estimated inbreeding coefficient. The Faroes region labels are: **VM** = Vágar and Mykines; **SR** = Suðuroy; **SM** = Suðurstreymoy; **SD** = Sandoy, Skúvoy, Stóra Dímun; **NG** = Norðoyggjar; and **EN** = Eysturoy og Norðstreymoy.

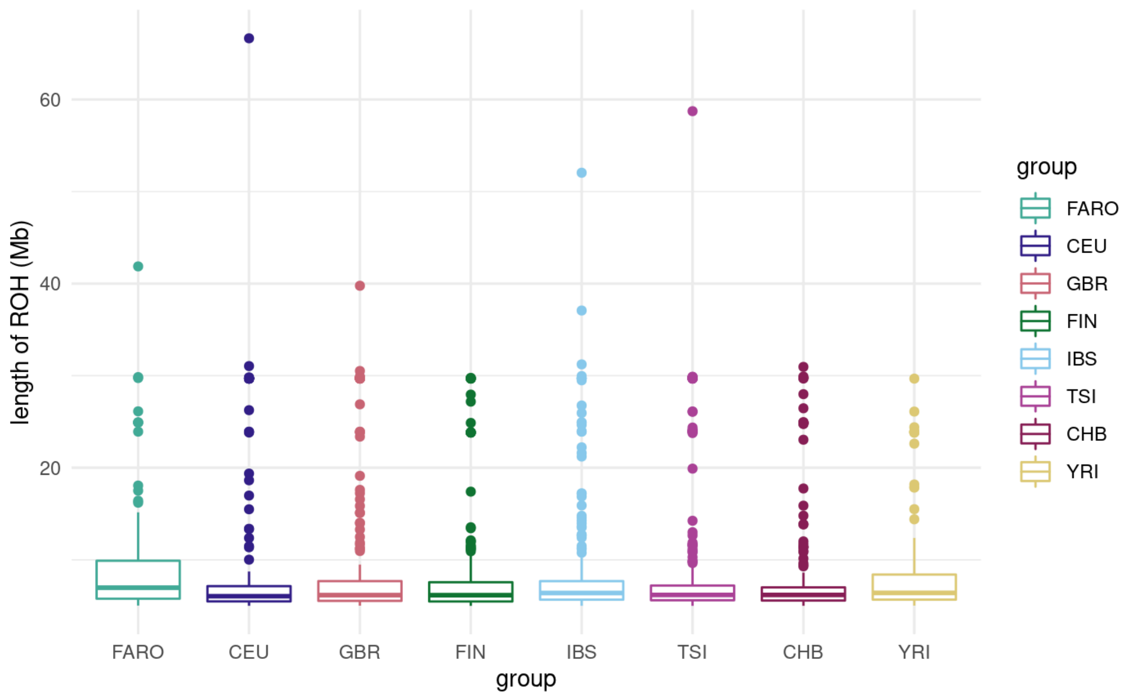


**Figure S3. ROH length distributions by group.** Length (Mb) of all runs of homozygosity (ROH) stratified by group.

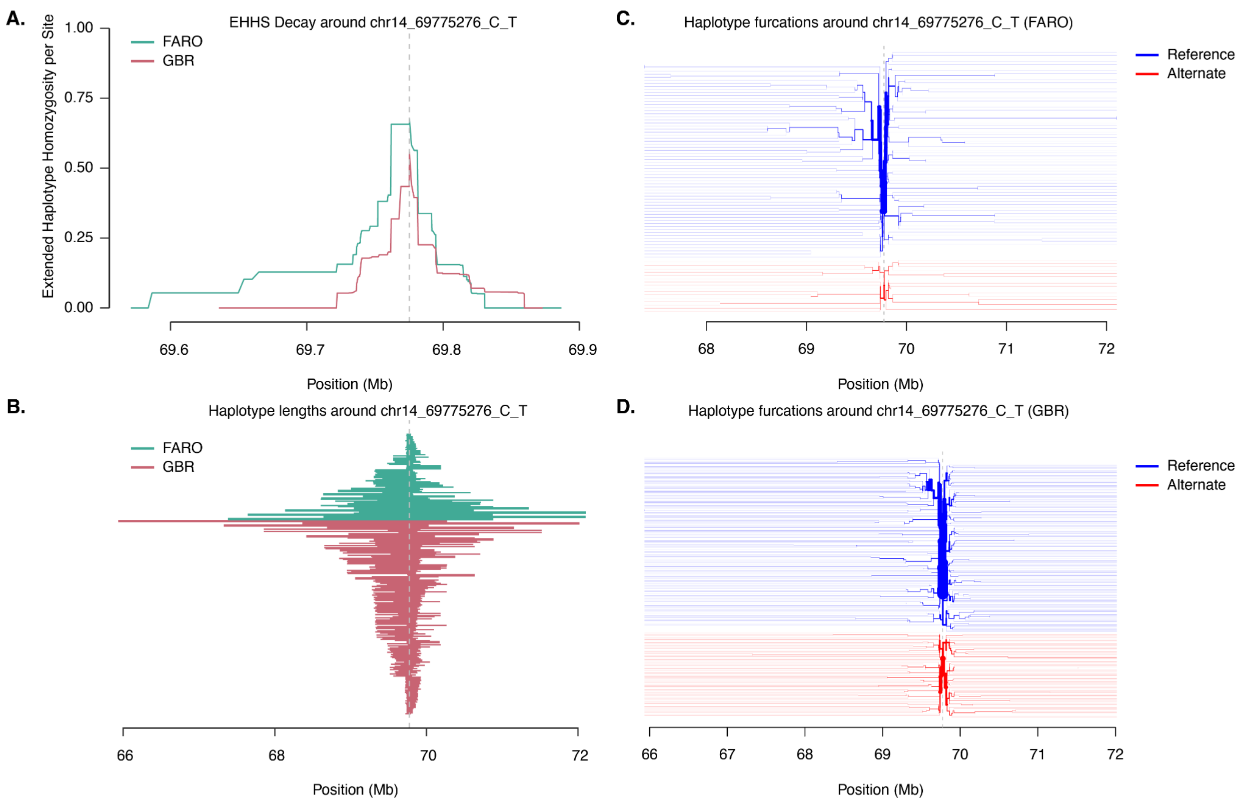


**Figure S4. Haplotype visualizations for top XP-EHH variant in *SLC10A1* / *SRSF5* locus. A)** Decay in Expected Haplotype Homozygosity per Site (EHHS) for chr14_69775276_C_T, comparing Faroese (FARO, teal) and British (GBR, red) haplotypes. **B)** Lengths for distinct haplotypes spanning chr14_69775276_C_T comparing FARO (teal) and GBR (red). **C)** haplotype furcation plot for FARO centered on chr14_69775276_C_T **D)** haplotype furcation for GBR centered on the same allele. In the haplotype furcation plots (panels **C** & **D**), haplotypes for the reference allele (C) are in blue, and those for the alternate allele (T) are in red.

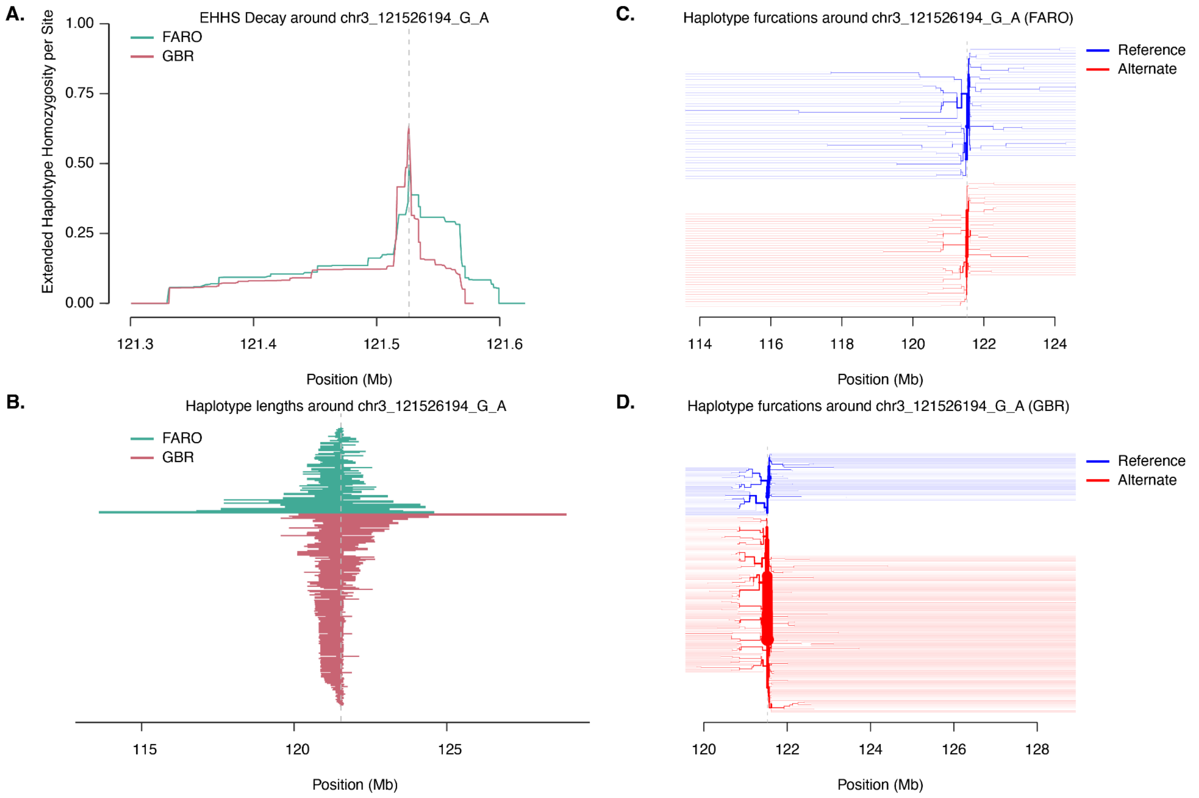


**Figure S5. Haplotype visualizations for top XP-EHH variant in *POLQ* locus. A)** Decay in Expected Haplotype Homozygosity per Site (EHHS) for chr3_121526194_G_A, comparing Faroese (FARO, teal) and British (GBR, red) haplotypes. **B)** Lengths for distinct haplotypes spanning chr3_121526194_G_A comparing FARO (teal) and GBR (red). **C)** haplotype furcation plot for FARO centered on chr3_121526194_G_A **D)** haplotype furcation for GBR centered on the same allele. In the haplotype furcation plots (panels **C** & **D**), haplotypes for the reference allele (G) are in blue, and those for the alternate allele (A) are in red.

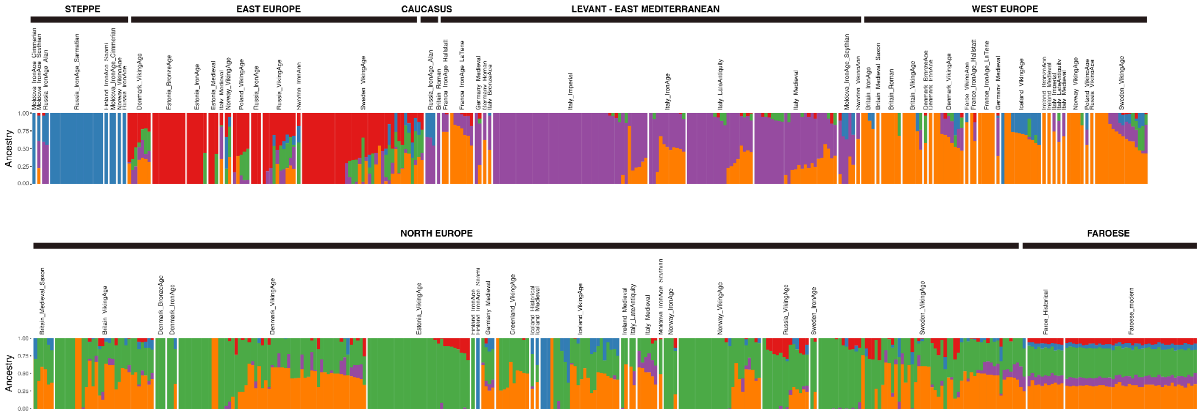


**Figure S6. Admixture plot showing proportions for 616 imputed ancient genomes from Europe together with 40 present-day Faroese genomes from this study.** Ancient individual groups are categorized based on patterns of IBD clustering as inferred in Allentoft *et al.* 2024. The plot uses five colors to represent different ancestral sources, which are maximized in individuals in different regions of europe: orange for “West Europe”, green for “North Europe”, blue for “Steppe”, purple for the “Levant and East Mediterranean” and red for “Eastern Europe”.

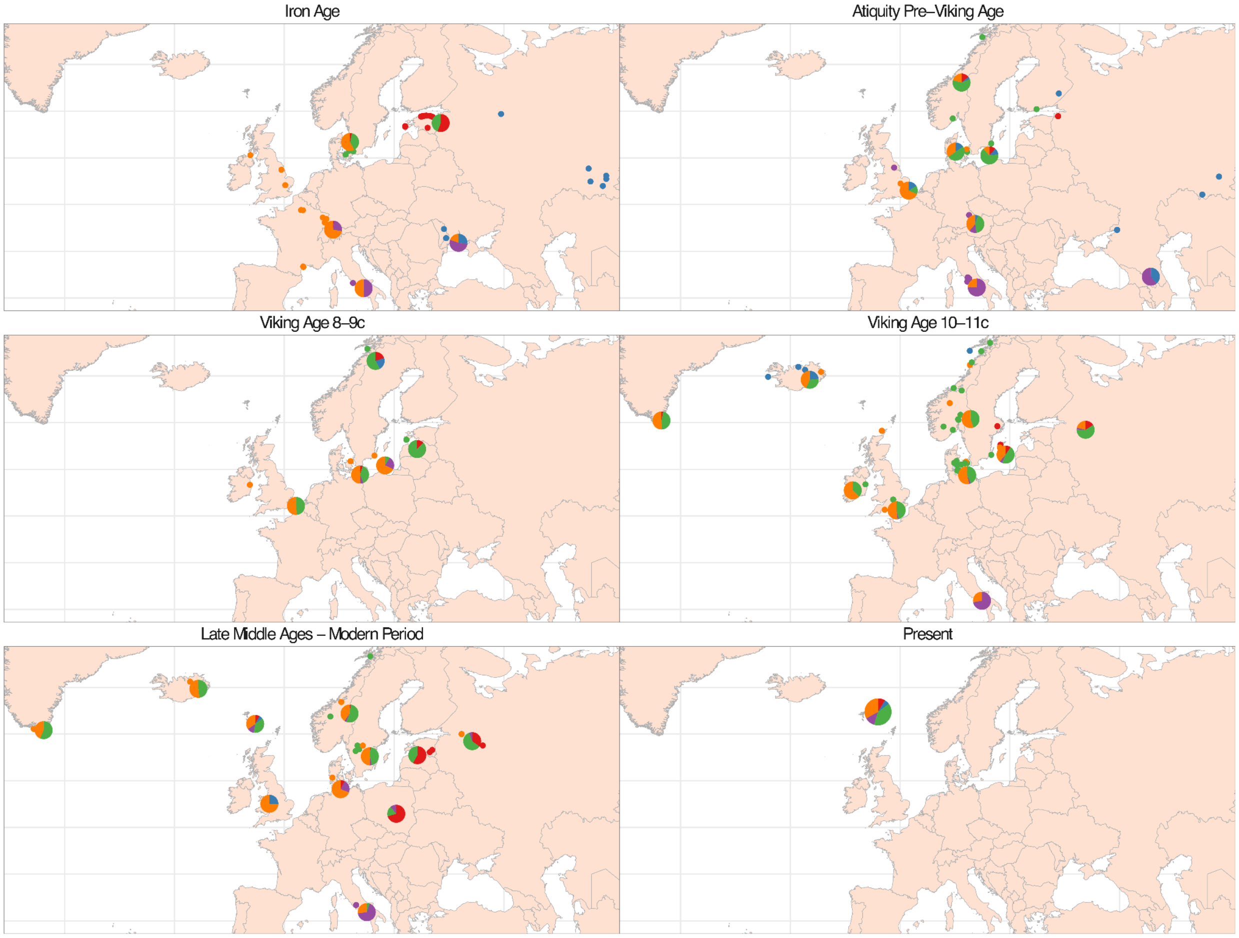


**Figure S7. Map illustrating the geographical distribution of ancestry proportions for 616 ancient imputed individuals from Europe and 40 modern Faroese genomes from this study.** The map is divided into panels to capture both geographical and temporal variations across Europe. Each pie chart on the map represents admixture proportions with five colors: green, orange, blue, red, and purple maximize Northern European, Celtic, Steppe, Eastern European, and Levant / East Mediterranean ancestries, respectively, reflecting different ancestral sources.

A.

B.

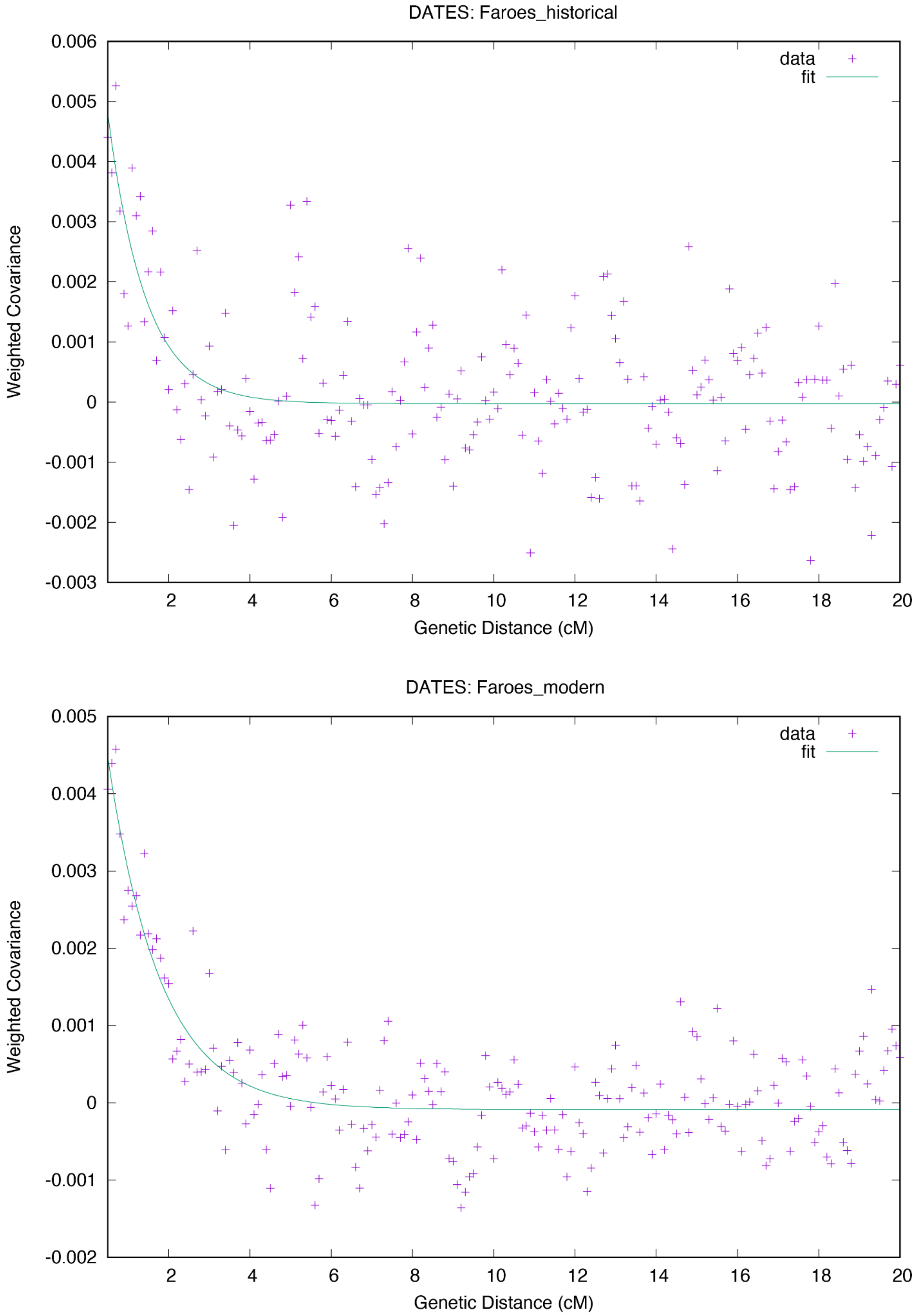


**Figure S8. *DATES* estimated ancestry covariance and least squares exponential fit.** Plots were output by the *DATES* software for **A)** 11 historical Faroese individuals (dated to approximately the 18th century) and **B)** 40 modern Faroese individuals.

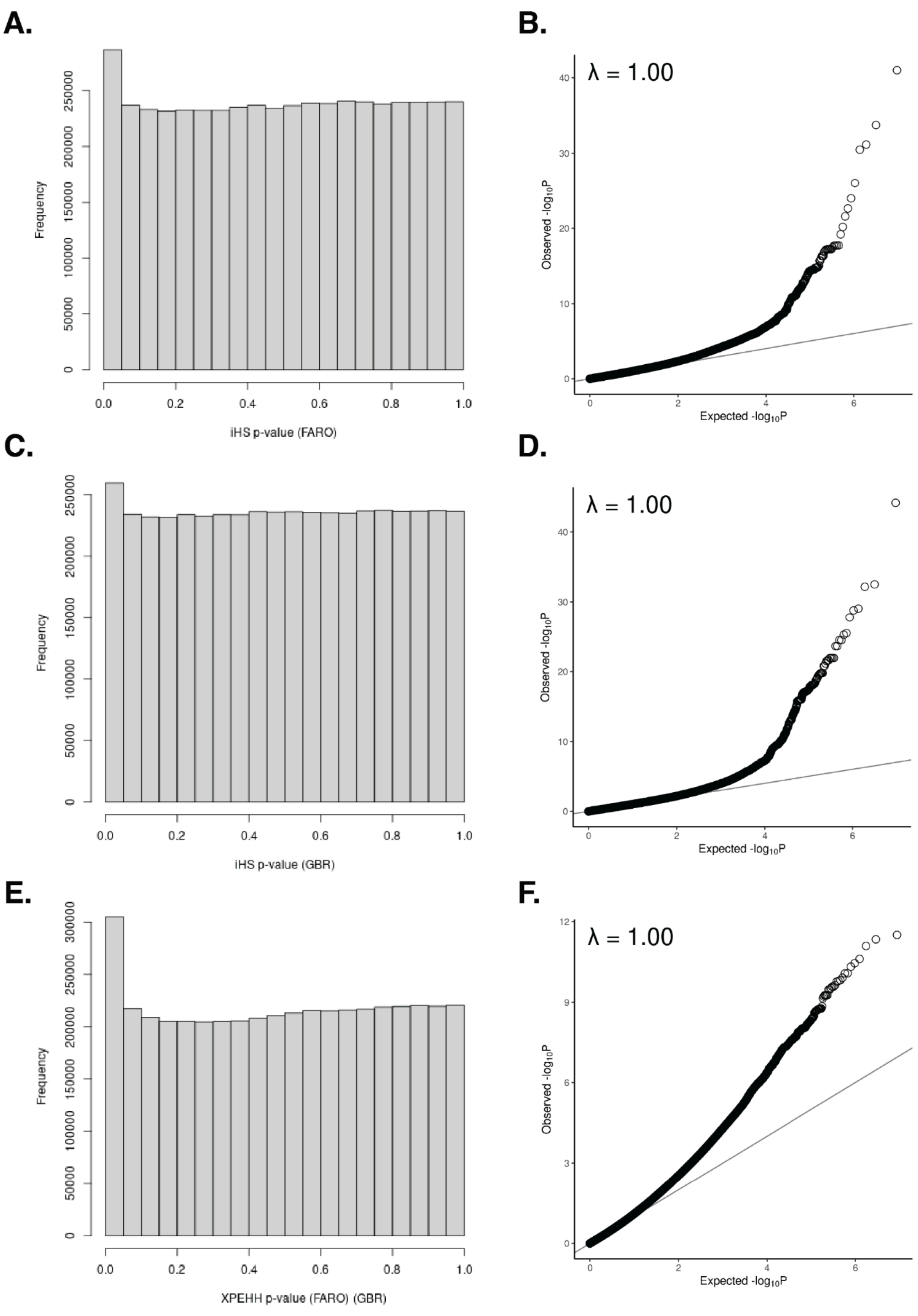


**Figure S9. Q-Q plots and p-value histograms for selection statistics.** Histograms of p-value distributions for **A)** integrated haplotype score (iHS) in the Faroese (FARO) haplotypes, **C)** iHS in the British (GBR) haplotypes, **E)** cross-population expected haplotype homozygosity (XP-EHH) comparing FARO and GBR. Q-Q plots for observed versus expected log transformed p-values for **B)** iHS in FARO, **D)** iHS in GBR **F)** XP-EHH between FARO and GBR. The estimated lambda inflation value is shown in each Q-Q plot.

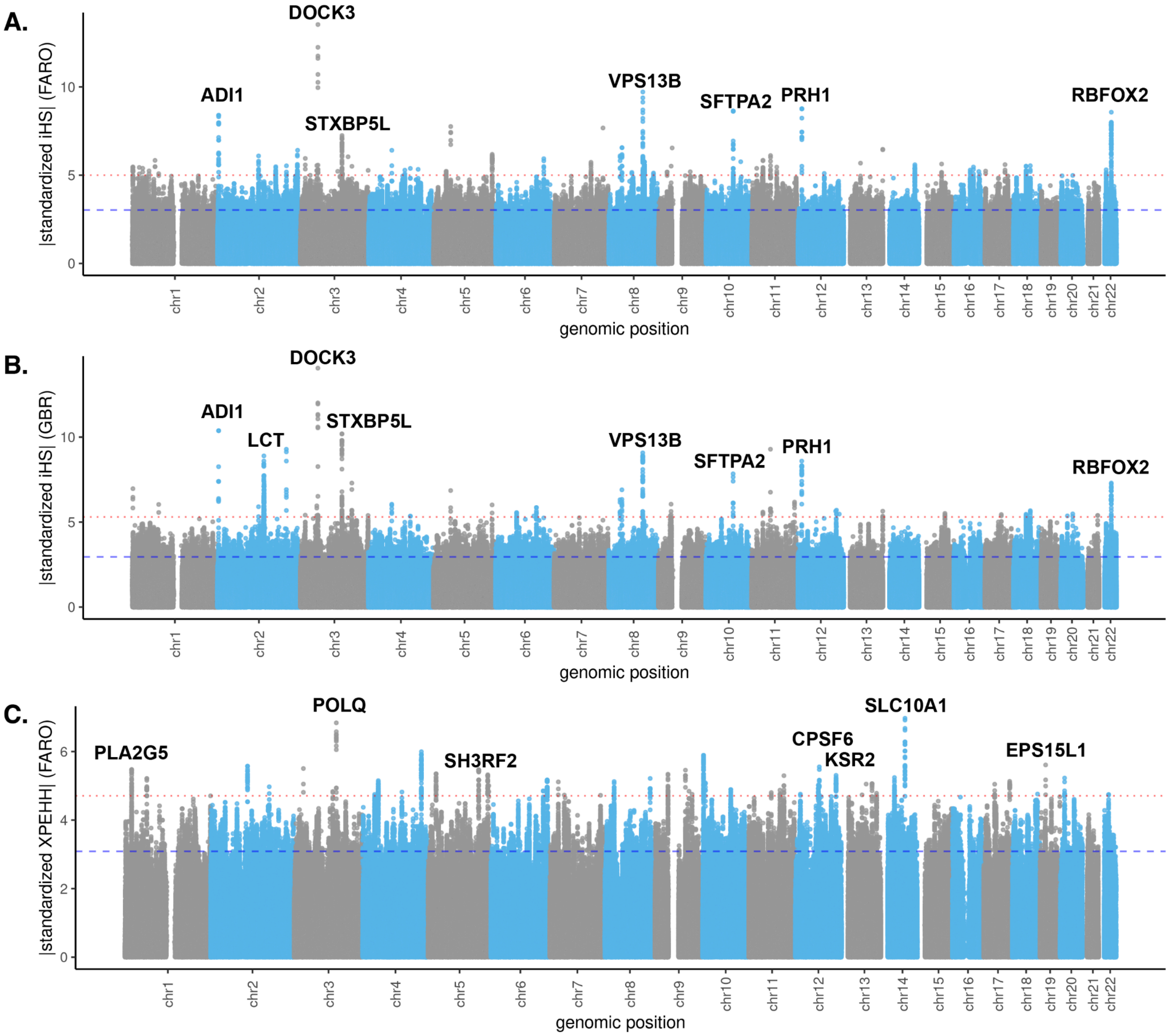


**Figure S10. Selection scan results for Faroese and British cohorts. A)** Absolute value of the standardized integrated haplotype score (iHS) in the 40 Faroese genomes (FARO). **B)** Absolute value of the standardized integrated haplotype score (iHS) for 90 British WGS samples from 1000 Genomes (GBR). **C)** The standardized cross-population expected haplotype homozygosity (XPEHH) for FARO (all positive values). The top 0.5% of results are indicated by the blue dashed line in each plot, and the top 0.01% of results are indicated by the red dotted line in each plot. Some genes in the top loci are indicated on each plot.
